# Identification of an intrinsically disordered region (IDR) in arginyltransferase 1 (ATE1)

**DOI:** 10.1101/2024.08.23.609426

**Authors:** Misti Cartwright, Rinky Parakra, Ayomide Oduwole, Fangliang Zhang, Daniel J. Deredge, Aaron T. Smith

## Abstract

Arginyltransferase 1 (ATE1) catalyzes arginylation, an important post-translational modification (PTM) in eukaryotes that plays a critical role in cellular homeostasis. The disruption of ATE1 function is implicated in mammalian neurodegenerative disorders and cardiovascular maldevelopment, while post-translational arginylation has also been linked to the activities of several important human viruses such as SARS-CoV-2 and HIV. Despite the known significance of ATE1 in mammalian cellular function, past biophysical studies of this enzyme have mainly focused on yeast ATE1, leaving the mechanism of arginylation in mammalian cells unclear. In this study, we sought to structurally and biophysically characterize mouse (*Mus musculus*) ATE1. Using size-exclusion chromatography (SEC), small angle X-ray scattering (SAXS), and hydrogen deuterium exchange mass spectrometry (HDX-MS), assisted by AlphaFold modeling, we found that mouse ATE1 is structurally more complex than yeast ATE1. Importantly, our data indicate the existence of an intrinsically disordered region (IDR) in all mouse ATE1 splice variants. However, comparative HDX-MS analyses show that yeast ATE1 does not have such an IDR, consistent with prior X-ray, cryo-EM, and SAXS analyses. Furthermore, bioinformatics approaches reveal that mammalian ATE1 sequences, as well as in a large majority of other eukaryotes, contain an IDR-like sequence positioned in proximity to the ATE1 GNAT active-site fold. Computational analysis suggests that the IDR likely facilitates the formation of the complex between ATE1 and tRNA^Arg^, adding a new complexity to ATE1 structure and providing new insights for future studies of ATE1 functions.

**For Table of Contents Use Only:** 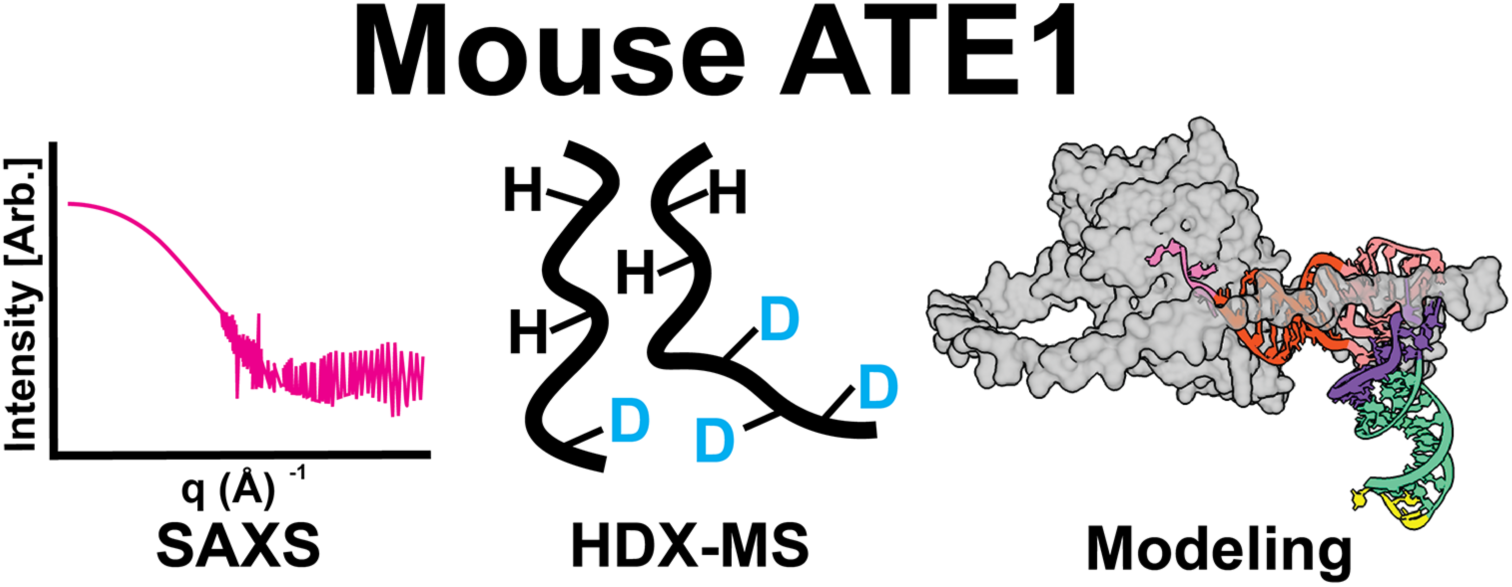

## INTRODUCTION

To increase the vastness of the proteome and to refine its dynamic nature, organisms alter polypeptides after protein translation by using post-translational modifications (PTMs),^1–4^ and an enigmatic yet essential PTM linked to normal eukaryotic cellular homeostasis is that of arginylation.^5–7^ Catalyzed by the enzyme arginyltransferase 1 (ATE1), arginylation is a transfer ribonucleic acid (tRNA)-dependent post-translational process in which the amino acid arginine (Arg) is transferred from the 3’ end of a donor tRNA to that of an acceptor protein (Fig. 1).^6,8,9^ Post-translational arginylation is essential to maintain cellular homeostasis in eukaryotes due to its involvement in several key biological processes. One such process is the N-degron pathway, a hierarchical pathway that links the identity of the N-terminal residue of a protein to its *in vivo* half-life. This is because arginylation can be added to the unprotected N-terminus of a peptide, either in freshly translated protein with the removal of the N-terminal methionine (Met) from the acceptor protein via Met aminopeptidase,^5,7^ or by protease-dependent internal polypeptide cleavage. ATE1 recognizes N-terminal negatively-charged amino acid sidechains (Glu, Asp, or oxidized Cys) and catalyzes the transfer of Arg from Arg-tRNA^Arg^ to the acceptor protein to form a new peptide bond in non-ribosomal manner.^6^ The N-terminally arginylated protein may be then recognized by N- recognins, ubiquitinylated, and degraded in a proteasomal-dependent manner.^7,10^ Examples of proteins N-terminally arginylated and subsequently degraded in this manner include regulators of G-protein signaling (RGS), such as RGS4, RGS5, RGS7, and RGS16 that are important in mammalian cardiovascular maturation as well as in the development of the mammalian nervous system.^6,11^ However, degradation is not the only fate for arginylated proteins, as this PTM may also alter the functions of the target proteins, commonly by causing changes in protein surface charges, often leading to changes in protein oligomerization. One example is that of the chaperone calreticulin, which dimerizes when it is arginylated leading to the formation of stress granules that increase cell survival in Ca^2+^-depleted conditions.^12^ Another example is the arginylation of the cytoskeletal protein β-actin, which is thought to play an important role in actin polymerization under certain biological contexts.^13,14^ In addition, research has shown that arginylation may also occur at a sidechain of an acidic amino acid (Glu or Asp) within the polypeptide (termed mid-chain arginylation) via formation of an isopeptide bond,^15^ although this type of modification is considered much less frequent. An example is that of the neural protein α-synuclein that may be stabilized in an arginylation-dependent manner, ameliorating the formation of plaques that contribute to neurodegenerative disease.^16^ These examples emphasize the importance of arginylation to eukaryotic function, but how this process occurs is unclear.

**Figure 1.**
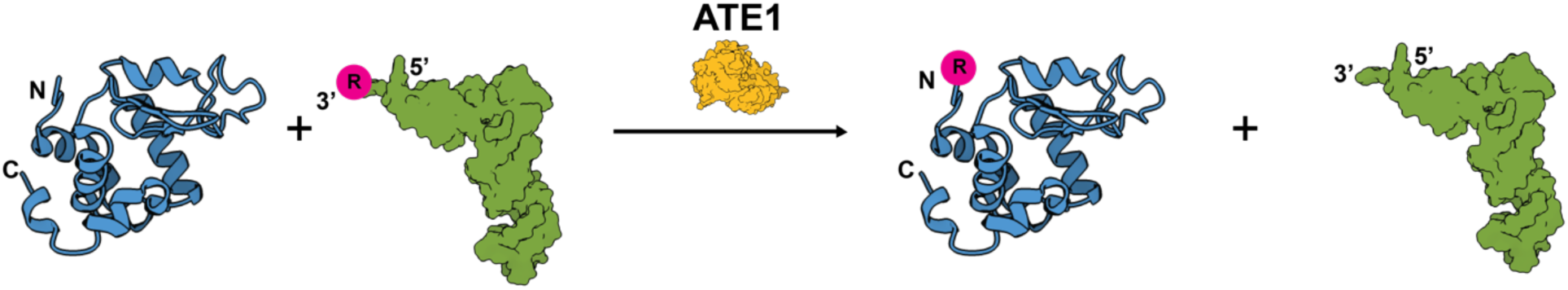
Cartoon depiction of eukaryotic post-translational arginylation. Catalyzed by the eukaryotic enzyme arginyltransferase 1 (ATE1; yellow), arginylation is the covalent and post-translational transfer of the amino acid arginine (R; pink) from the 3’ end of an aminoacylated tRNA (green) to an acceptor protein (blue). Arginylation occurs most commonly at the N-terminus of target polypeptides bearing negatively-charged sidechains (such as Asp, Glu, and oxidized Cys). Arginylation plays many important degradative and non-degradative biological roles in eukaryotic cellular homeostasis. The labels ‘N’ and ‘C’ represent the locations of the N- and C-termini, respectively.

The last two years have witnessed a major leap forward in deciphering the structural and regulatory mechanisms of post-translational arginylation. In 2022, the X-ray crystal structures of ATE1 from two yeast species (*S. cerevisiae* and *K. lactis*) were determined (Fig. 2 a,b),^17,18^ revealing the overall three-dimensional fold of ATE1 for the first time. The structure of yeast ATE1 reveals a bilobed protein fold containing a GCN5-related *N*-acetyltransferase (GNAT) domain known to be the location of catalytic activity (Fig. 2). N-terminally adjacent to the GNAT fold is a small domain that is Cys-rich and binds an [Fe-S] cluster that regulates arginylation activity both *in vitro* and *in viv*o.^19,20^ However, under certain conditions, a Zn^2+^ ion is also capable of occupying the [Fe-S] site,^17^ but its functional relevance is unknown. In 2023, a cryo-EM structure of yeast ATE1 bound to bacterial tRNA^Arg^ was determined. This structure revealed that the acceptor stem of the tRNA only makes modest surface contacts with the GNAT fold (Fig. 2c), but the 3’ end of the tRNA buries into the GNAT pocket, presumably to maintain the key electrostatic interactions between the conserved GNAT Asp/Glu residue and the Arg covalently linked to the tRNA (Fig. 2c). These observations are consistent with prior hypotheses, ^17,18,21,22^ and these exciting results have provided evidence for a proposed molecular mechanism of post-translational arginylation.^17,18,22^ However, while the study of yeast ATE1 has been important, a major gap remains in understanding the mechanism of arginylation in mammalian cells, which is critical due to ATE1’s important connection to various disease-like states in humans,^23–27^ and without which rational design of therapeutics is inaccessible. However, While yeast ATE1 and most mammalian ATE1s (such as mouse ATE1) are virtually identical in size (*ca.* 59 kDa monomer vs. *ca*. 60 kDa monomer, yeast and mouse ATE1, respectively), they have significant difference in their amino acid sequence (generally <30 % sequence similarity) and other properties such as predicted isoelectric points (predicted pI ≈ 5.8 vs. 8.3, yeast and mouse ATE1, respectively), creating a barrier for applying the structural insights obtained from yeast ATE1 to mammalian ATE1. Interestingly, mammalian ATE1 sequences have regions of high charge and low complexity, suggesting that an intrinsically disordered region (IDR) may be present, distinct from the structures of yeast ATE1s.

**Figure 2.**
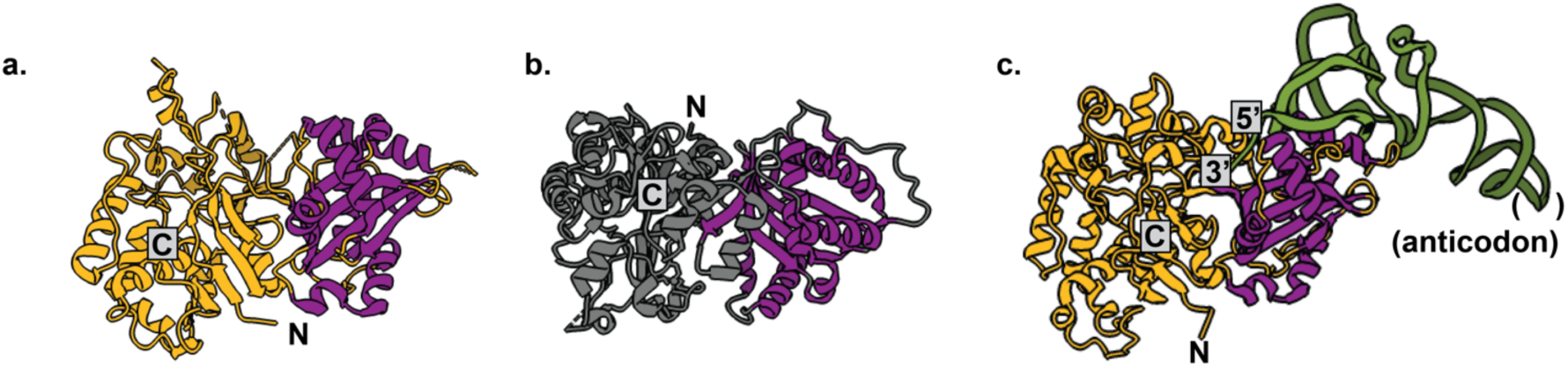
Recent structural advancements in eukaryotic post-translational arginylation. **a**. X-ray crystal structure of *S. cerevisiae* ATE1 (PDB ID 7TIF) with the conserved active site GNAT fold highlighted in purple. **b**. X-ray crystal structure of *K. lactis* ATE1 (PDB ID 7WFX) with the conserved active site GNAT fold highlighted in purple. **c**. Cryo-EM structure of *Sc*ATE1 in complex with tRNA^Arg^ (PDB ID 8FZR), with the tRNA colored in green and the conserved active site GNAT fold highlighted in purple. The labels ‘N’ and ‘C’ represent the locations of the N- and C-termini, respectively.

IDRs are domains of polypeptides with low complexity that are known to be important for numerous cellular functions such as division, transcription, and even translation.^28,29^ In general, a protein IDR may be superficially characterized by a span of amino acids with low hydrophobicity but high overall net charge, which often prevents the formation of an otherwise globular or partially globular domain (although small micro domains may be folded in IDRs).^30,31^ More than simply unfolded, however, these regions of polypeptides are generally intrinsically flexible and highly dynamic, which cannot be assigned by protein sequence or even structural modeling alone.^31,32^ IDRs may function as flexible linkers to allow for protein domains to be independently folded and to function separately.^33^ Importantly, IDRs may also assist in the formation of macromolecular interactions, such as in the organization of protein-nucleic acid complexes (more common) or protein-protein complexes (less common).^30,34^ These interactions may even nucleate protein and/or nucleic acid folding.^35,36^ While the significance of IDRs in the functions of proteins, particularly the nucleotide-interacting proteins are well demonstrated, their existence in ATE1, a tRNA-binding protein, remained unknown.

In this work, we use a suite of computational and biophysical approaches to understand the structure of mammalian ATE1. We initially model *Mus musculus* ATE1 isoform 1 (*Mm*ATE1-1) using the AlphaFold server, which predicts a core bi-lobed structure containing a similar GNAT fold like yeast ATE1 but connected via a large, unstructured linker of random coil that does not exist in yeast ATE1. To verify the accuracy of the AlphaFold model, we then express and purify recombinant forms of *Mm*ATE1-1 for crystallization trials and biophysical characterization. We use small-angle X-ray scattering (SAXS) and find that *Mm*ATE1-1 is chiefly monomeric in solution, similar to yeast ATE1, and mostly globular with exception of the large, unstructured linker region that is predicted to be highly flexible in solution based on molecular dynamics simulations. Importantly, using hydrogen-deuterium exchange mass-spectrometry (HDX-MS), we then demonstrate that this linker in *Mm*ATE1-1 is indeed an IDR, and we confirm that such a structure does not exist in yeast ATE1s. Further sequence analyses indicate that a similar IDR may exist in many, if not most, higher-order ATE1s. Finally, computational modeling suggests that the ATE1 IDR may have evolved for purposes of increasing tRNA-protein interactions, which is likely to affect ATE1 function. When taken all together, this study provides insight into the mechanism of mammalian arginylation and suggests that most ATE1 proteins contain an IDR, which has not been shown previously and dramatically changes our understanding of the structure of ATE1.

## RESULTS

### AlphaFold modeling suggests MmATE1-1 contains the core bilobed ATE1 fold but with a large, unstructured linker region

We initially used AlphaFold to model the structure of *Mus musculus* (mouse) ATE1 isoform 1 (*Mm*ATE1-1), which we previously studied in the context of [Fe-S] cluster binding.^20^ Intriguingly, modeling predicts similarities to, but also important differences from, the experimentally-determined structures of yeast ATE1s. As seen in Fig. 3b, the AlphaFold model of *Mm*ATE1-1 is predicted to have a core folded region containing the N-terminal regulatory domain (comprising residues *ca*. 1-79 including the CxxC and CC motifs) and a C-terminal domain (comprising residues *ca*. 407 to 516) of unknown function folded together and directly adjacent to the enzymatic GNAT fold that is split (comprising residues *ca*. 80-103 and *ca*. 253-406). Superpositioning of the predicted AlphaFold *Mm*ATE1-1 GNAT domain with that of the experimentally-determined *Sc*ATE1 GNAT domain predicts high structural homology in the enzymatic active site (C_α_ RMSD *ca*. 0.8 Å across the entire GNAT fold; Fig. 3c), and the key sidechain carboxylate buried in the hydrophilic GNAT α-β-α sandwich that is believed to make electrostatic contacts with the 3’ Arg on Arg-tRNA^Arg^ aligns nearly perfectly between *Sc*ATE1 (Asp277) and *Mm*ATE1-1 (Glu385) (Fig. 3d). Notably in the *Mm*ATE1-1 AlphaFold model, however, is the presence of a large, unstructured region of high charge and low hydrophobicity that splits the GNAT fold and is inserted between residues *ca*. 104-252 (Fig. 3a.e). The confidence of the AlphaFold model in this region is generally low, but this unstructured region is predicted to first feed *ca*. 30 amino acids through the cleft at the lobe-lobe interface between the N-terminal regulatory domain and the GNAT fold before extending unstructured into space prior to linking back up with the GNAT fold (Fig. 3e). Importantly, this region of the GNAT fold encompasses both the ATE1 tRNA-binding site in yeast ATE1, as well as the hypothesized substrate-binding site of the ATE1 arginylation target protein. Using a sequence alignment, this linker region appears to have been an insertion at some point in evolution and is not present in the sequence of *Sc*ATE1 (Fig. S1) nor in its X-ray or cryo-EM structures (Fig. 2b). To determine whether the unstructured linker region was a product of the various splicing-derived isoforms of *Mm*ATE1 that may be produced, we also used AlphaFold to predict the structures of splice variants *Mm*ATE1-2 to *Mm*ATE1-4, and each displayed the presence of a similar large, unstructured region of random coil splitting the GNAT fold (Figs. S2, S3), consistent with this region lacking variability from isoform to isoform (Fig. S2). However, as these are merely predicted structures, we then sought to determine experimental support for the AlphaFold model of *Mm*ATE1-1.

**Figure 3.**
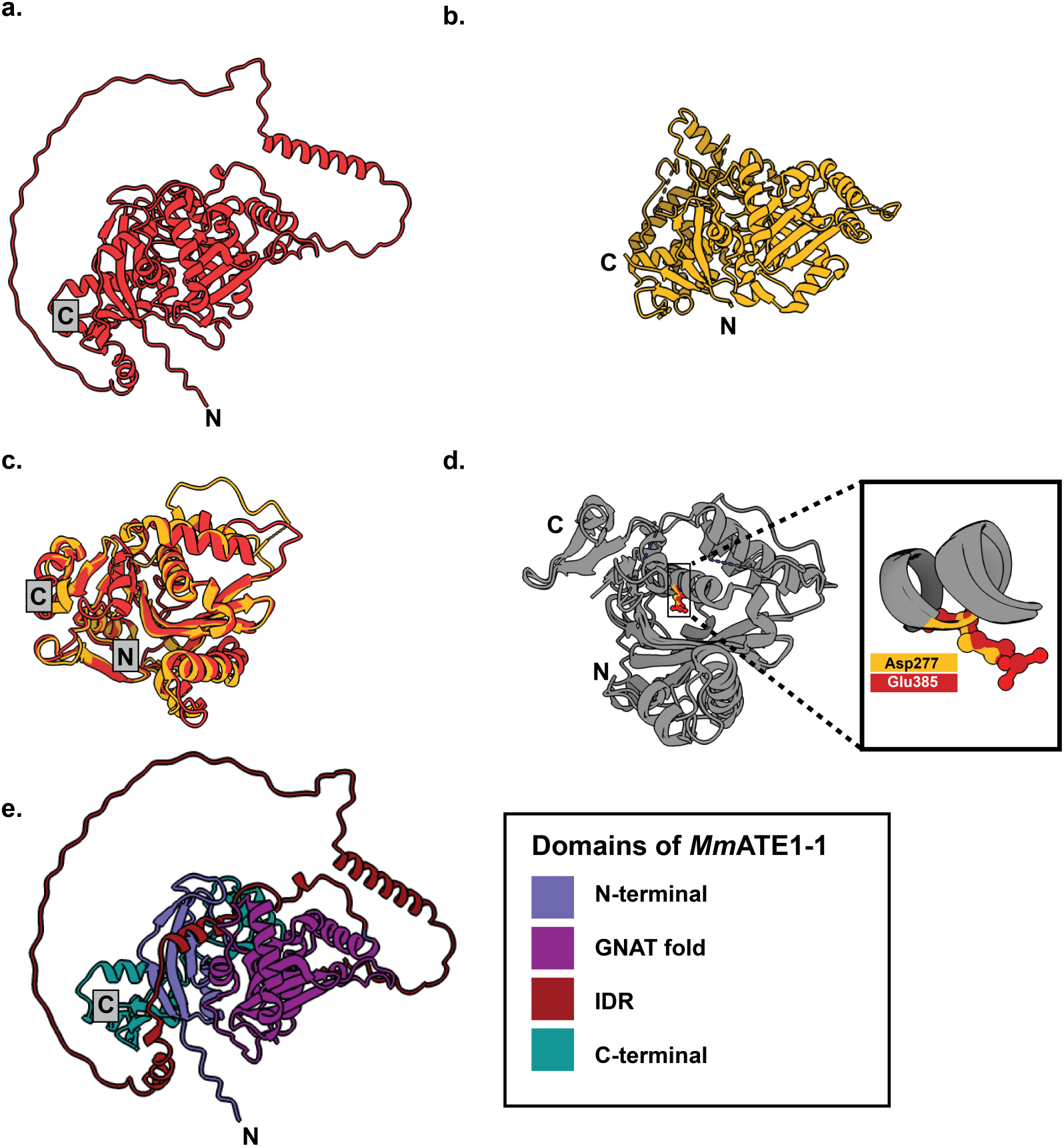
Computational modeling suggests that mouse ATE1 is more complex than yeast ATE1. **a**. AlphaFold model of *Mm*ATE1-1. **b**. X-ray crystal structure of *Sc*ATE1 (PDB ID 7TIF). **c**. Superpositioning of the AlphaFold *Mm*ATE1-1 GNAT domain with that of the *Sc*ATE1 GNAT domain suggests structural conservation of the core enzymatic GNAT fold. **d**. Zoomed-in view of the key amino acid (Asp277 in *Sc*ATE1, yellow; Glu385 in *Mm*ATE1-1, red) proposed to make electrostatic interactions with Arg on the charged Arg tRNA^Arg^. **e**. The various domains of the AlphaFold *Mm*ATE1-1 model color coded: N-terminal (royal blue); GNAT fold (purple); IDR (brick red); C-terminal (teal). The labels ‘N’ and ‘C’ represent the locations of the N- and C-termini, respectively.

### Recombinantly expressed MmATE1-1 is highly homogeneous and monomeric

To build upon our previous successes in solving the X-ray crystal structure of *S. cerevisiae* (yeast) ATE1,^18^ and to provide support for the *Mm*ATE1 AlphaFold models, we then sought to express and purify a mammalian ATE1 to determine its structure, which was achieved with the similar protocol in our past study to obtain enzymatically active *Mm*ATE1, splice variant1 (*Mm*ATE1-1).^20^ Consistent with our prior *Sc*ATE1 results,^20^ *Mm*ATE1-1 oxically purified through immobilized metal affinity chromatography (IMAC) and then size-exclusion chromatography (SEC) was extensively brown in color and bore the hallmarks of a partially degraded [Fe-S] cluster known to bind to the N-terminal regulatory domain of ATE1. To simplify handling of the protein for crystallization trials, we removed the [Fe-S] cluster from the protein by ethylenediaminetetraacetic acid (EDTA) treatment while simultaneously cleaving the C-terminal (His)_6_ tag with Tobacco Etch Virus (TEV) protease. In both cases, regardless of whether the tag was cleaved or the tag was intact, the isolated protein was highly pure and appeared to be mainly monomeric (Fig. S4), like that of *Sc*ATE1.^37^ However, we were unable to obtain any crystals of *Mm*ATE1-1 (cleaved or uncleaved forms) when utilizing similar conditions to that of *Sc*ATE1 (successfully crystalized) or by setting up extensive sparse-matrix crystallization trials. Importantly, difficulty in crystallizing proteins with IDRs is not uncommon because these flexible, dynamic domains often preclude the formation of a stable crystalline lattice. Furthermore, because the predicted IDR domain of *Mm*ATE1 is located internal to the polypeptide (Fig. 3), it could not be easily removed unlike a flexible N- or C-terminus. Therefore, we turned to alternative methods to characterize *Mm*ATE1-1 structurally.

### SAXS supports the predicted MmATE1-1 AlphaFold model and suggests the large, unstructured linker region is highly dynamic

To understand the gross structural features and dynamics of *Mm*ATE1-1, we sought to use small-angle X-ray scattering (SAXS) to characterize this protein in solution and to compare its in-solution behavior to that of *Sc*ATE1.^18^ To determine whether any differences in the SAXS profile might occur, we generated both the cleaved and uncleaved apo forms of *Mm*ATE1-1 and collected high-quality size-exclusion chromatography (SEC)-coupled SAXS data on these proteins (Fig. 4a). For both cases, concentrations from 2-8 mg/mL of each protein were tested, and multi-angle light scattering (MALS) performed in tandem with SEC-SAXS showed that both protein constructs were generally monomeric in solution, although a modest shoulder suggesting the presence of a minor amount of dimeric protein was seen at higher concentrations (estimated at <20 % of the total protein), consistent with our gel filtration experiments (*vide supra*), but distinct from human ATE1.^37^ A Guinier analysis of both constructs at 9.0 mg/mL (intact) and 6.5 mg/mL (tag-cleaved) *Mm*ATE1-1 showed strong linearity at high q values (Fig. 4b) indicating minimal aggregation was present in the analyzed samples and allowing us to estimate the radius of gyration (*R_g_*) for both proteins: 29.5 Å for uncleaved *Mm*ATE1-1 and 28.3 Å for cleaved *Mm*ATE1-1 (Table 1). In both samples, the Kratky analysis suggests the presence of a mostly globular protein based on the shapes of the curve ranging from *ca*. 0-4 qR_g_ (Fig. 4c), but in the extended region from *ca*. 5-10 qR_g_, the shape of the curve suggests an additional domain with flexibility may be present.^38,39^ As this observation is present regardless of the presence/absence of the C-terminal affinity tag, the Kratky analysis suggests the majority of flexibility lies elsewhere in the protein. In contrast, the pairwise distance distribution function (*P(r)*) analysis on both forms of *Mm*ATE1-1 suggests modest differences in the maximum dimension of the particle in solution (*D*_max_) in solution: 115Å for uncleaved *Mm*ATE1-1 and 98 Å for cleaved *Mm*ATE1-1, suggesting that the affinity tag may contribute modestly to differences in the observed SAXS data (Table 1). Interestingly, both the R_g_ and *D*_max_ values for *Mm*ATE1-1 are larger than the respective values for *Sc*ATE1 (27.9 Å and 82 Å, respectively), despite the similar molecular weights of the two proteins, and the Kratky analysis of *Mm*ATE1-1 indicates that this protein may be generally more flexible than *Sc*ATE1, which may be a result of the unstructured domain observed in the AlphaFold model of *Mm*ATE1-1 but absent in the structure of *Sc*ATE1 (Fig. 3).

**Figure 4.**
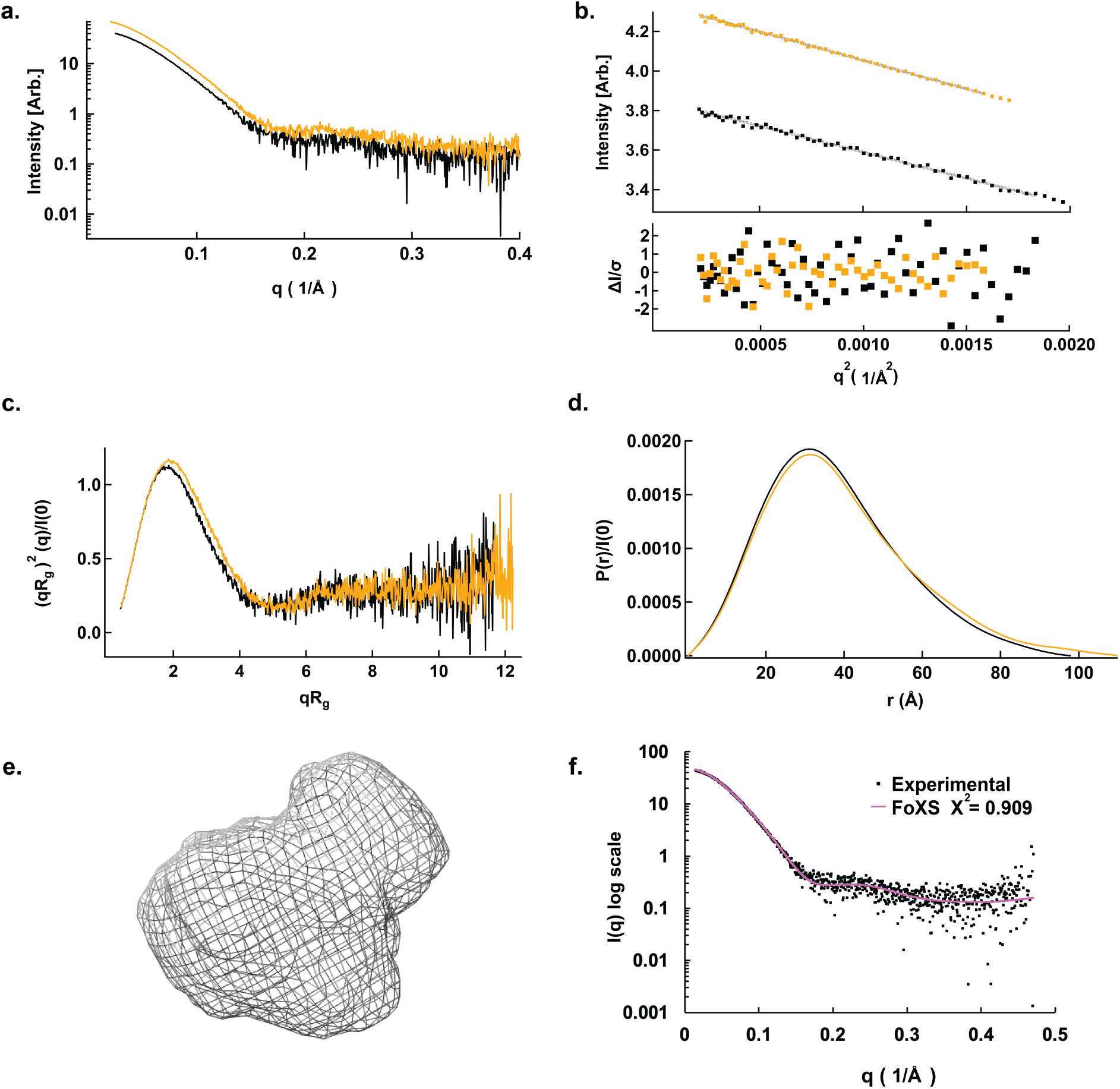
SAXS analyses of apo *Mm*ATE1-1 with the affinity tag cleaved (black) and uncleaved (yellow). **a**. The log10 plot scattering profiles of cleaved and uncleaved *Mm*ATE1-1. **b**. The Guinier region of *Mm*ATE1-1 cleaved and uncleaved with their linear fits (gray; top panel), and the respective residues from the fits (bottom panel). **c**. The Kratky plots of cleaved and uncleaved *Mm*ATE1-1 calculated from the experimental SAXS data. **d**. The *P(r)* distribution plots of cleaved and uncleaved *Mm*ATE1-1. **e**. The *ab initio* envelope (gray mesh) generated from the SEC-SAXS data of cleaved *Mm*ATE1-1. **f**. The experimental SAXS data of cleaved *Mm*ATE1-1 (black squares) overlaid with the FoXS-calculated SAXS profile (pink) generated from the BilboMD simulations. The simulations generate good agreement based on the calculated *χ*^2^ value of 0.909.

**Table 1.** SAXS data table, including information regarding the Guinier analysis, the *P*(*r*) analysis, and DAMAVER calculations for cleaved and uncleaved *Mm*ATE1-1.

| **Guinier analysis** |  | **Cleaved *MmATE1-1*** | **Uncleaved *MmATE1-1*** |
| --- | --- | --- | --- |
| $I(0)$ [Arb.] | | $47.33 \pm 0.13$ | $77.02 \pm 0.22$ |
| $R_g$ (Å) | | $28.28 \pm 0.11$ | $29.50 \pm 0.21$ |
| $q_{\min}$ (Å<sup>-1</sup>) | | 0.265 | 0.471 |
| $q_{\max}R$ | | 1.311 | 1.323 |
| Coefficient of correlation, $R^2$ | | 0.963 | 0.999 |
| ***P(r)* analysis** |  |  |  |
| $I(0)$ (cm<sup>-1</sup>) | | $47.77 \pm 0.13$ | $77.80 \pm 0.16$ |
| $R_g$ (Å) | | $28.93 \pm 0.10$ | $30.47 \pm 0.11$ |
| $D_{\max}$ (Å) | | 98 | 115 |
| $q$ range (Å<sup>-1</sup>) | | 0.021 - 0.44 | 0.021 - 0.41 |
| $\chi^2$ (total estimate from GNOM) | | 0.759 | 0.909 |
| **DAMAVAR** |  |  |  |
| Symmetry, anisotropy assumptions |  | P1, unknown | P1, unknown |
| Models averaged (omitted) |  | 15 (0) | 15 (0) |
| Mean NSD (S.D.) | | $1.30 \pm 0.12$ | $1.29 \pm 0.16$ |
| Resolution (Å) |  | 38 | 38 |

We then compared the *Mm*ATE1-1 AlphaFold model to the *Mm*ATE1-1 SAXS data in order to understand whether the predicted model and experimentally-determined data were in agreement. Using DAMMIN and DMAVER,^40,41^ we first generated an *ab initio* envelope for *Mm*ATE1-1 both with and without the affinity tag, which were both very similar, but we chose to focus on the cleaved *Mm*ATE1-1 to preclude any contributions that the presence of the affinity tag may have caused. In general, the *ab initio* enveloped of cleaved *Mm*ATE1-1 (Fig. 4e) suggests the presence of two features, one larger and one smaller, in the low-resolution reconstruction, distinct from that of *Sc*ATE1 that is mostly globular.^18^ The AlphaFold model of *Mm*ATE1-1 nicely resembles the *ab initio* envelope generated from the cleaved *Mm*ATE1-1 SAXS data (Fig. S5) with the exception of the large, unstructured linker region. Reasoning that the large, unstructured linker region may not fit well into the *ab initio* envelope due to its potential flexibility, we then used BilboMD to generate an ensemble of conformations of the *Mm*ATE1-1 AlphaFold model (Fig. S6).^42^ Indeed, this ensemble displays dramatic conformational changes within the large, unstructured linker region that, when averaged, nicely recapitulate (𝜒^2^ *ca*. 0.9) the experimentally-determined *Mm*ATE1-1 SAXS data (Fig. 4f).^43^ Combined, these data not only suggest that the core domain of *Mm*ATE1-1 is similar to the AlphaFold model, but also that the large, unstructured linker region is likely highly dynamic, indicating that this part of the polypeptide behaves like an intrinsically-disordered region (IDR).

### HDX-MS identifies the presence of an intrinsically-disordered region in MmATE1

To demonstrate that the large, unstructured linker region in the *Mm*ATE1-1 model is indeed intrinsically disordered in solution, we then used hydrogen-deuterium exchange mass-spectrometry (HDX-MS). To do so, we measured the deuterium uptake of *Mm*ATE1-1 as a function of time (10 s, 100 s, and 1000 s) along with a fully-deuterated (FD) control. After deuteration, digestion, and mass spectrometry analyses, a total of 124 peptides were obtained corresponding to a sequence coverage of 97.4% of the primary sequence of the *Mm*ATE1-1 protein with an average redundancy of 3.25 (Fig. 5a,b, Fig. S7). The HDX-MS Woods plot of *Mm*ATE1-1 shows rapid and high relative fractional uptake corresponding to a central region of the *Mm*ATE1-1 protein (Fig. 5a, Fig. S8). We then mapped these results onto the AlphaFold model of *Mm*ATE1-1 with highly dynamic residues colored in red while less dynamic residues are colored in blue (Fig. 5b). Consistent with our hypothesis, the region of strongest relative fractional deuterium uptake aligns almost perfectly to our theorized IDR (Fig. 5b). This dramatic relative fractional uptake of deuterium is highly rapid and complete typically within 1-2 min of deuterium incubation (Fig. 5c), indicating that the large linker region in the *Mm*ATE1-1 model is largely unstructured, consistent with our initial hypothesis from the AlphaFold model and our molecular dynamics-coupled SAXS analyses (*vide supra*) that the linker region is intrinsically disordered. For instance, peptides 141-168 and 178-201(Fig. 5c, I-II) show a relative deuterium uptake of 100% at all time points with very little variation suggesting a highly labile region of the protein. Peptides 213-228 (Fig. 5c, III), which correspond to the alpha helix in the IDR region are slightly less labile, perhaps due to the helical secondary structure. When the peptides in the IDR region are compared to other regions such as the peptides 441-450 (Fig. 5c, IV) we see that there is a more progressive increase in deuteration as a function of time showing that this region is structured compared to those in the IDR. HDX-MS supports that the rest of the *Mm*ATE1-1 protein is generally folded in solution and less solvent accessible than the IDR, with the exception of the first 11 residues at the N-terminus (predicted random coil in the AlphaFold model), a portion of the C-terminal domain (residues *ca*. 494-510) that may be dynamic in solution, and the C-terminal affinity tag when present (Fig. 5b, Fig S9). Thus, we can conclude from our HDX-MS data that the region of residues *ca*. 104-252 in the *Mm*ATE1-1 model indeed comprises an IDR.

**Figure 5.**
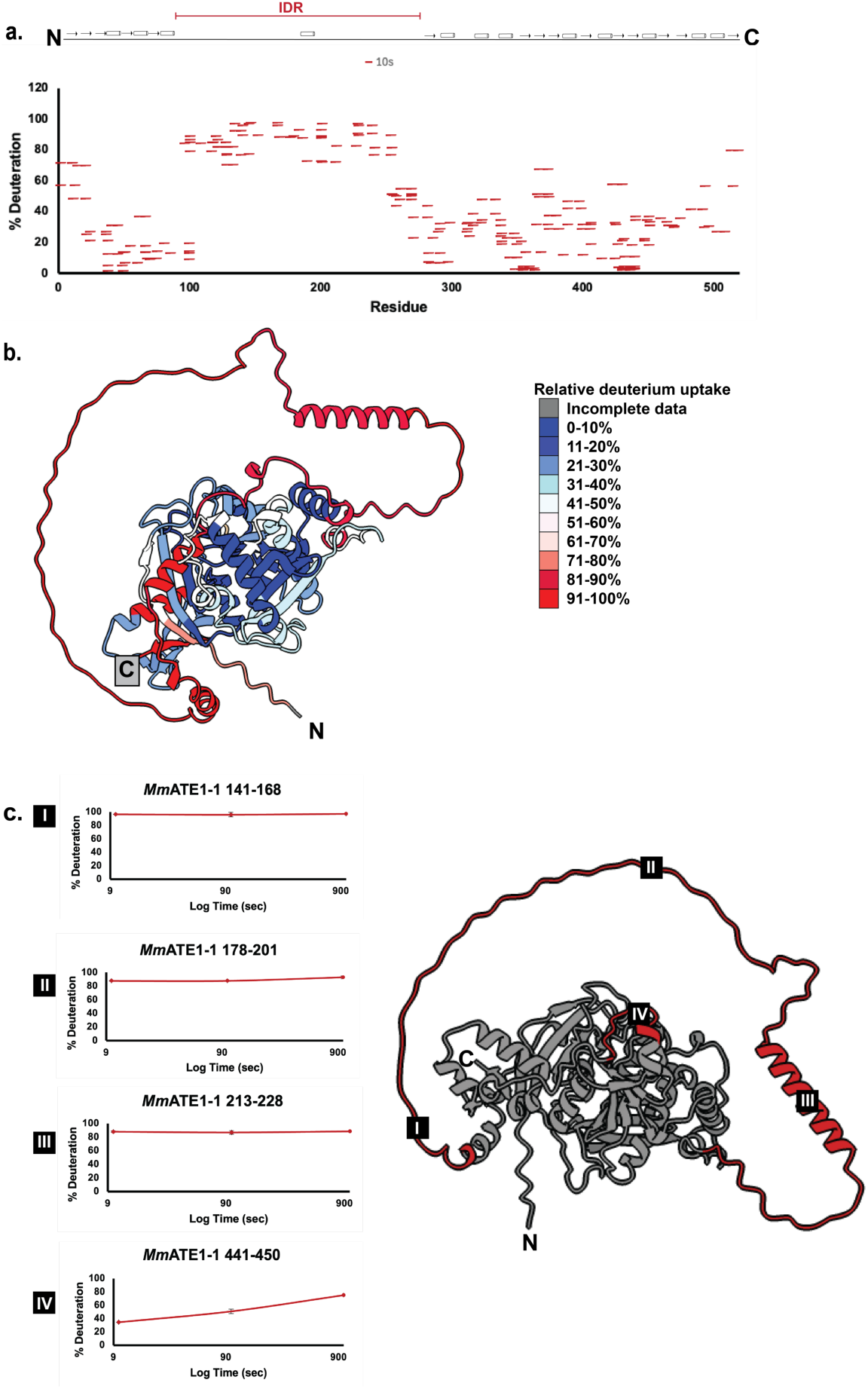
HDX-MS data on *Mm*ATE1-1. **a**. Woods plot of *Mm*ATE1 representing percent deuteration values at 10s in the apo state, plotted against amino acid sequence. Each bar represents a unique *Mm*ATE1 peptide that had a significant deuterium uptake at 10s. A cartoon of the *Mm*ATE1-1 polypeptide is represented at the top, drawn from N- to C-terminus, and the region corresponding to the IDR is labeled in red. **b**. Relative deuterium uptake of *Mm*ATE1-1, color coded from blue (minimal uptake) to red (maximal uptake) overlaid on the *Mm*ATE1-1 AlphaFold model. Gray labeling represents areas without data in the FD control. **c**. Kinetic deuteration plots of select *Mm*ATE1-1 peptides from areas of the IDR (labeled I, II, and III) and a disordered region near the GNAT fold (IV). The relative locations of these peptides are labeled on the AlphaFold model of *Mm*ATE1-1. The labels ‘N’ and ‘C’ represent the locations of the N- and C-termini, respectively.

### An IDR is likely present in the majority of ATE1 enzymes but not in yeast ATE1

Considering the known significance of IDRs in protein structure and function, we then sought to use bioinformatics to understand whether an IDR may be a general structural feature of ATE1 or if the IDR were only present in a few select species. First, using ATE1 sequences from a variety of organisms across the tree of life, we were able to generate a multiple sequence alignment (MSA) and qualitatively look for the presence of an insertion within the N-lobe of ATE1 that splits the GNAT fold (Fig. S10). In general, the presence of the IDR insertion within the GNAT fold appears to be present in all isoforms of higher-order eukaryotes such as fly, plant, mouse, and even human ATE1; however, even some ostensibly simpler eukaryotes such as slime mold are also predicted to have the IDR insertion. We then used a phylogenetics analysis to see if a more robust pattern would arise that could give insight into the point in evolution where the IDR was inserted into the GNAT fold (Fig. S11). While the phylogenetics analysis recapitulates the general trend observed in the MSA, no clear bifurcation appears in which a divergence can be attributed to the presence of an IDR in ATE1 (Fig. S11). In general, beyond complex fungi such as *Rhizopus stolonifer* and *Mitosporidium daphniae*, there appears to be an IDR-like region in the ATE1 amino acid sequence, suggesting that the IDR insertion may be present in the majority of ATE1s (Figs. S10, S11).

A notable exception to this generalization is that yeast ATE1s (such as *S. cerevisiae* and *K. lactis*) do not appear to have an IDR based on a lack of sequence conservation, which we then validated using HDX-MS. To do so, we repeated our previous experiments this time on apo *Sc*ATE1 (previously structurally characterized using both X-ray crystallography and cryo-EM). We measured deuterium uptake of *Sc*ATE1 over time (10 s, 100 s, and 1000 s) and compared the behavior to fully deuterated controls After deuteration, digestion, and mass spectrometry analyses, we were able to obtain a total of 56 peptides corresponding to a final sequence coverage of 81.9 % of the primary sequence of the *Sc*ATE1 with a redundancy of 2.0 (Fig. S12). In general, in *Sc*ATE1 we do not observe a low-complexity region of the size and the extent of that in *Mm*ATE1-1, indicating that yeast ATE1 does not have an IDR (Fig. 6a,b, Figs S13, S14), which is also consistent with the MSA analysis (Figs. S1, S10). However, there are some smaller regions of the yeast protein that may be highly dynamic in solution that are likely critical to function, such as: the region from *ca*. 128-139 in *Sc*ATE1 (capping the GNAT fold in the X-ray crystal structure and making contact with tRNA in the cryo-EM structure), the region from *ca*. 302-312 in *Sc*ATE1(at the lobe-lobe interface and unable to be visualized in the X-ray crystal structure), the region from *ca*. 335-375 in *Sc*ATE1 (a region of random coil folded against the body of *Sc*ATE1), and the C-terminal affinity purification tag (Fig. 6c). When taken together, these results suggest that the bulk of ATE1 sequences have an IDR region that interacts with the ATE1 GNAT fold, but notably not yeast ATE1s, which have been extensively characterized using both X-ray crystallography and cryo-EM.

**Figure 6.**
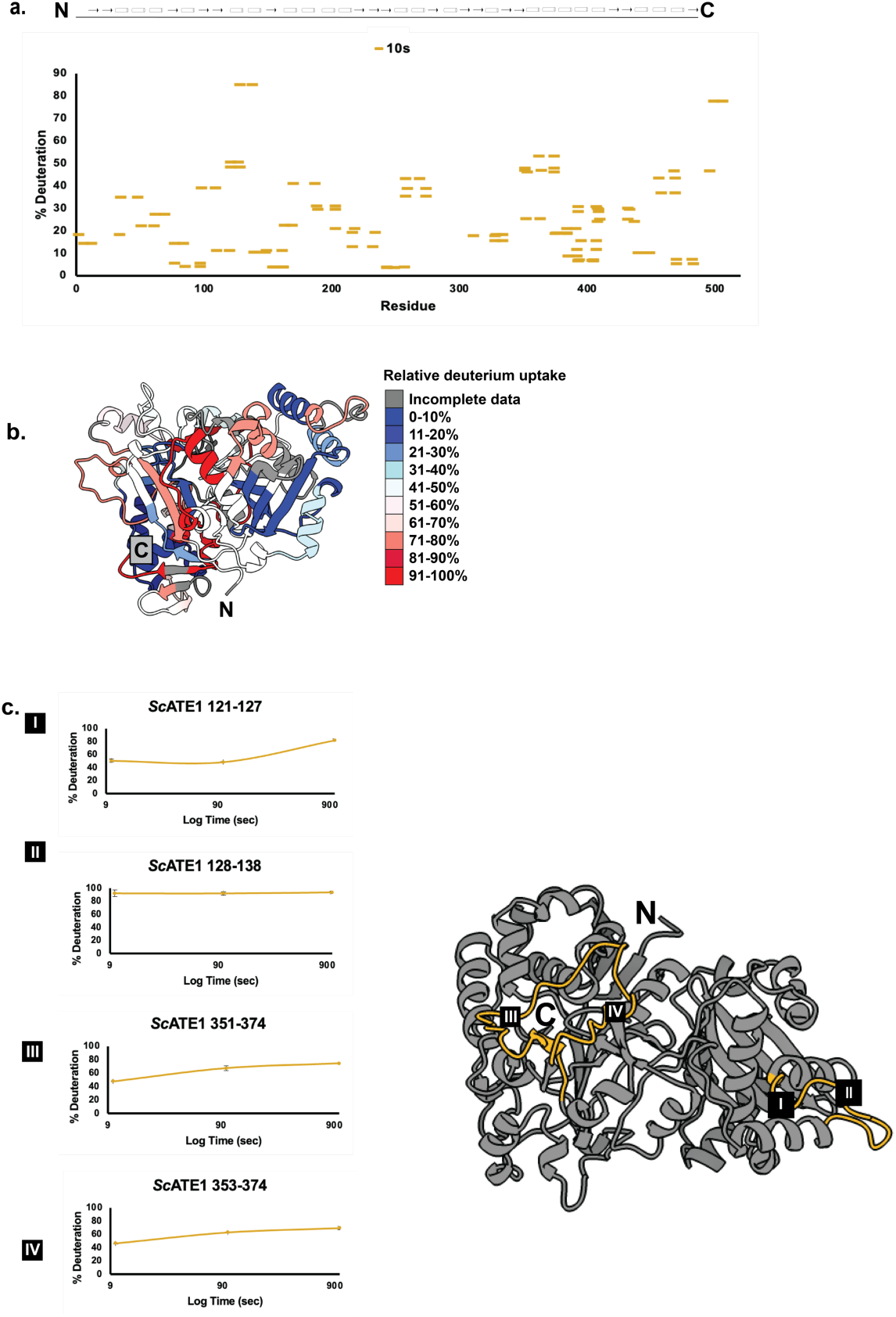
HDX-MS data on *Sc*ATE1. **a**. Woods plot of *Sc*ATE1 representing percent deuteration values at 10 s in the apo state, plotted against amino acid sequence. Each bar represents a unique *Sc*ATE1 peptide that had a significant deuterium uptake at 10s. A cartoon of the *Sc*ATE1 polypeptide is represented at the top, drawn from N- to C-terminus. **b**. Relative deuterium uptake of *Sc*ATE1, color coded from blue (minimal uptake) to red (maximal uptake) overlaid on the *Sc*ATE1 structure. Gray labeling represents areas without data. **c**. Kinetic deuteration plots of select *Sc*ATE1 peptides from areas of the *Sc*ATE1 structure (labeled I, II, III, and IV). While there are some regions that are disordered in solution based on the HDX-MS analysis, there is no large IDR present in *Sc*ATE1 like there is in *Mm*ATE1. The labels ‘N’ and ‘C’ represent the locations of the N- and C-termini, respectively.

### The IDR may be involved in ATE1-tRNA interactions and have a regulatory function

Given our new knowledge that *Mm*ATE1-1 has an IDR that is positioned within proximity of the tRNA-binding site along the ATE1 GNAT fold, we hypothesized that one role of the IDR could be to interact with tRNA in order to enhance tRNA binding to ATE1. It is known that only the acceptor stem of tRNA is required for arginylation, representing only a modest portion of the tRNA macromolecule, but how ATE1 effectively binds (and selects) tRNA is still unclear.^21,22^ As our SAXS and HDX-MS data suggested the AlphaFold modeling of apo *Mm*ATE1-1 was highly accurate, we then tested AlphaFold modeling of *Mm*ATE1-1 with an Arg-specific tRNA^Arg^ molecule, as ATE1 is known to bind uncharged tRNA^Arg^.^22^ Intriguingly, the AlphaFold model of this complex suggests that the ATE1 IDR region loops around the tRNA molecule, likely to enhance its interactions with ATE1 (Fig. 7a,b). While the relative positioning of the anticodon and the anticodon loop of the tRNA are predicted to be variable, AlphaFold modeling consistently positions approximately 75 amino acids of the IDR to “wrap” around both the acceptor stem region, large areas of the T-loop, and small area of the tRNA D-loop (Fig. 7c). Additionally, this interaction between the ATE1 IDR and the tRNA is also predicted to prevent the IDR from sampling binding within the GNAT fold cleft (Fig. 7b compared to Fig. S6) where it would otherwise block entry of the protein substrate to the ATE1 GNAT active site. Thus, modeling suggests that the IDR we have newly identified in this work could be important for ATE1-tRNA interactions (consistent with concurrent observations in press on human ATE1),^37^ and that this IDR may also serve a regulatory function to control ATE1 activity.

**Figure 7.**
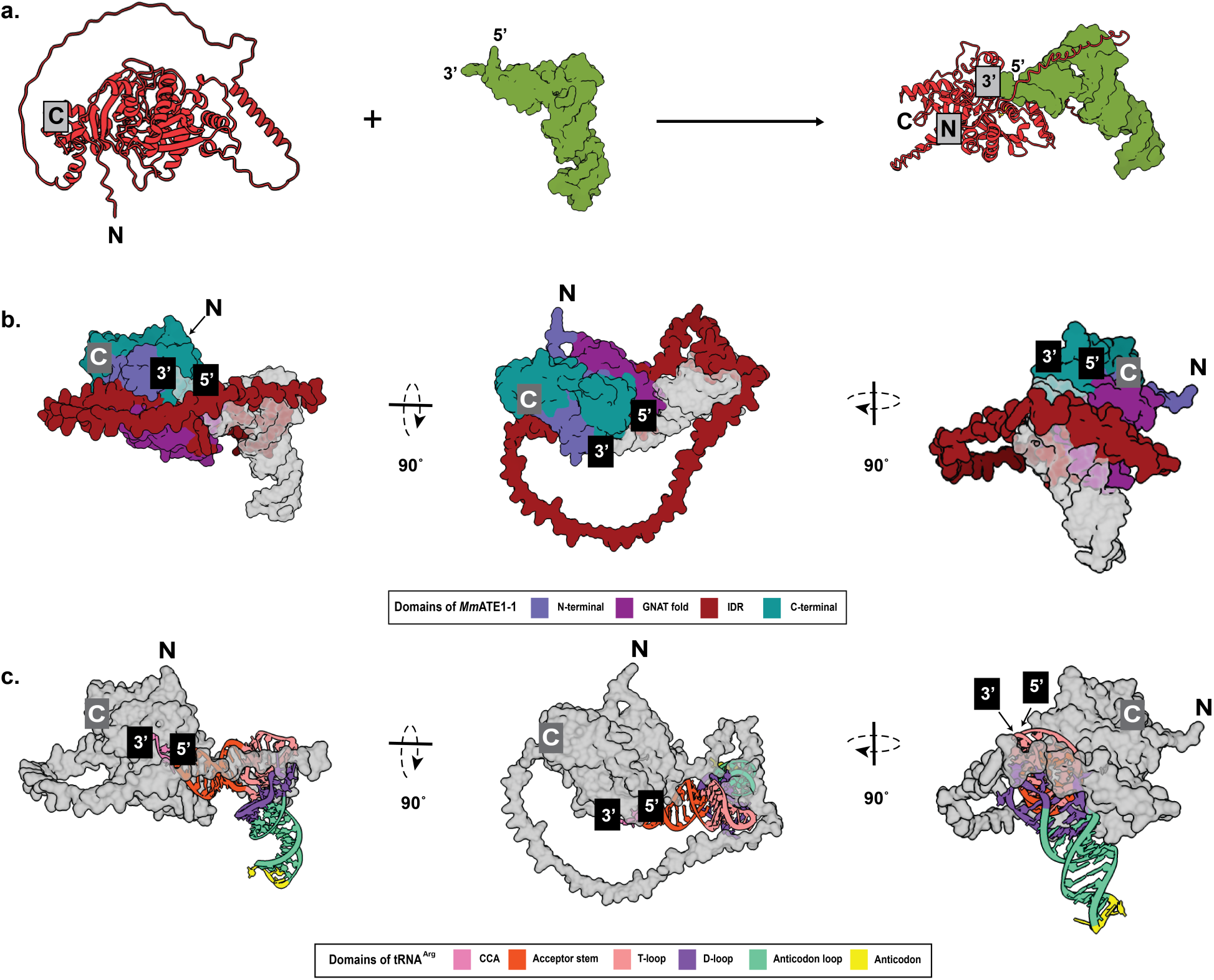
Computational modeling predicts a role for the ATE1 IDR. **a**. The AlphaFold model of *Mm*ATE1-1 (red) bound to tRNA^Arg^ (green) suggests that the ATE1 IDR makes extensive interactions with the tRNA acceptor stem and T-loop, likely enhancing the affinity between the two macromolecules. **b**. The predicted *Mm*ATE1-1-tRNA^Arg^ complex with domains of *Mm*ATE1-1 color-coded the same as in Fig. 3e, illustrating extensive contacts between the IDR and the tRNA. **c**. The predicted *Mm*ATE1-1-tRNA^Arg^ complex with regions of the tRNA color-coded, illustrating that the ATE1 IDR makes extensive contacts with the acceptor stem and the T-loop of the tRNA and minor contacts with the tRNA D-loop. In all panels, the labels ‘N’ and ‘C’ represent the locations of the N- and C- termini, respectively, and the 3’ and 5’ regions of the tRNA are also labeled.

## DISCUSSION

In this work, we have used multiple biophysical techniques to identify and to characterize the mammalian ATE1 IDR, which may be a common feature to most ATE1 enzymes. The presence of a large, unstructured domain with low hydrophobicity but high overall net charge (hallmarks of an IDR) was initially suggested based on AlphaFold modeling and sequence analyses of all *Mm*ATE1 isoforms. We then sought to purify mouse ATE1 for crystallization attempts, but we were unable to generate crystals of this enzyme despite its similarity in size to its yeast counterparts, likely due to the presence of this highly dynamic and unstructured region, requiring us to use alternative biophysical approaches. We then analyzed apo *Mm*ATE1-1 using SEC-coupled SAXS, and we found that this ATE1 isoform behaves monomerically in solution regardless of the presence or absence of the purification tag, similar to yeast ATE1 but distinct from the recently described behavior of human ATE1.^37^ Further analyses of the mouse ATE1 SAXS profile showed that this enzyme is both more flexible and generally larger in solution than yeast ATE1. We then coupled our AlphaFold modeling with molecular dynamics to demonstrate that the large, unstructured linker in the mouse ATE1 model was highly dynamic in solution, and the inclusion of this largely dynamic region could faithfully recapitulate the experimental solution SAXS data of *Mm*ATE1-1. To confirm that this region on *Mm*ATE1-1 is indeed intrinsically disordered, we used HDX-MS and showed rapid deuterium uptake that could be readily mapped onto the IDR within the AlphaFold model. This behavior contrasted strongly with the behavior of yeast ATE1 analyzed in solution under identical conditions, indicating that yeast ATE1 lacks an IDR. Indeed, multiple sequence alignments show that this region is conspicuously absent in yeast ATE1s that have been structurally characterized previously,^17,18,22^ but the IDR appears to be conserved in many mammalian ATE1 sequences, including human ATE1, changing our understanding of the general structure of ATE1 in an important way.

Based on this work, our data suggest that the ATE1 IDR may have two critical roles in the mechanism of arginylation, which could be potentially targeted for therapeutic developments. As modeled in Fig. 7, *ca*. 75 amino acids of the IDR in mouse ATE1 are predicted to “wrap” around a portion of the acceptor stem as well as the T-loop of tRNA^Arg^, making significantly more contact between ATE1 and the tRNA than observed in the previously published *Sc*ATE1-tRNA complex.^18^ While this model of the mouse ATE1-tRNA^Arg^ complex is computational, it is consistent with a manuscript just in press detailing the cryo-EM structure of human ATE1 bound to uncharged tRNA^Arg^.^37^ While only a loop of the human ATE1 IDR could be modeled into the cryo-EM map of human ATE1-tRNA^Arg^ complex, the region of the human ATE1 IDR that was observed makes contact between human ATE1 and the T-loop of the tRNA^Arg^,^37^ similar to our model of mouse ATE1-tRNA^Arg^ presented here. While it is not possible to easily remove the IDR from the ATE1 polypeptide, mutations at the human ATE1-tRNA^Arg^ interface weakened the binding affinities with the uncharged tRNA^Arg^ as estimated by electrophoretic mobility shift assays (EMSAs),^37^ supporting our hypothesis here that the IDR is important in increasing ATE1-tRNA affinity. However, unobserved in the aforementioned human ATE1 structure is the rest of the IDR, which is substantial and is predicted in our MD simulations to thread *ca*. 30 amino acids into the lobe-lobe interface, directly adjacent to the mouse ATE1 enzyme active site. Interestingly, this region includes Asp117 (*Mm*ATE1-1 and -2)/Asp110 (*Mm*ATE1-3 and -4), which is a known site of self-arginylation in three of the four major mouse ATE1 isoforms.^23^ It is tempting to speculate that this region of the IDR may partially occupy or occlude the enzyme active site until ATE1 undergoes self-arginylation, at which point the dynamics of the IDR could shift to allow for the binding of the substrate polypeptide. Of course, we cannot rule out additional possibilities for the role of the ATE1 IDR, such as its potential involvement in the formation of the proposed post-translational arginylation super complex,^44^ or its potential interactions with the ligand of ATE1 (Liat1), also known to be intrinsically disordered and to shuttle ATE1 into the nucleus of cells under certain conditions.^45,46^ In all of these scenarios, the IDR could play an integral role in ATE1 function, which might be targeted for therapeutic developments. While generating small molecules that select for an IDR may be complex due to its lack of defined tertiary structure, there have been recent advancements.^47–49^ Moreover, the ATE1 IDR appears to mediate tRNA interactions with the well-folded GNAT domain, which could provide an additional pocket for small-molecule binding.

The results from this work also provide insights into future structure-function studies of ATE1. As this work demonstrates that the ATE1 IDR is both substantial in size and highly dynamic in solution, this newly-discovered domain may limit modeling of this region in the electron density of future ATE1 X-ray crystal structures, or it may preclude the crystallization of additional ATE1s entirely, at least in their apo form. Based on our computational modeling, the presence of tRNA and/or substrate may attenuate the dynamicism of the ATE1 IDR, which could suggest that crystallization of these proteins is more likely to occur in complex with the tRNA macromolecule. As another option, isolated domains (such as the GNAT fold with a substantially truncated IDR) could be crystallized in the presence of substrates and, potentially, tRNA fragments that lack the IDR-interacting T-loop and D-loop. However, our work here suggests that there is likely strong structural homology between the core catalytic domain of yeast and mammalian ATE1s, and details on the domains flanking the core GNAT fold are critical to understanding the arginylation mechanism. Alternatively, cryo-EM has emerged as a viable approach for ATE1 structural studies,^18,22^ and at least part of the ATE1 IDR is visible in the very recent cryo-EM structure of human ATE1,^37^ but only when bound to tRNA. Based on our studies here, it is likely that a suite of complementary methods (such as SAXS, HDX-MS, X-ray crystallography, and cryo-EM) will be necessary to understand the structure and the dynamics of the mammalian arginylation complex fully, especially for the design of targeted therapeutics. Future exploration will work to uncover this exciting new complexity in ATE1 function.

## Acknowledgements

This work was supported by NIH-NIGMS grant R35 GM133497 (A.T.S), NIH-NIGMS grant T32 GM066706 (A. T. S. and M. C.), G-RISE grant T32-GM144876, NIH-NIGMS R01 GM138557 (F. Z.), and in part by the University of Maryland Baltimore, School of Pharmacy Mass Spectrometry Center (SOP1841-IQB2014). SAXS experiments were conducted at the Advanced Light Source (ALS), a national user facility operated by Lawrence Berkeley National Laboratory on behalf of the Department of Energy, Office of Basic Energy Sciences, through the Integrated Diffraction Analysis Technologies (IDAT) program, supported by the DOE Office of Biological and Environmental Research. Additional support comes from the National Institute of Health project ALS-ENABLE (P30 GM124169) and a High-End Instrumentation Grant S10OD018483. Sequence searches utilized both database and analysis functions of the Universal Protein Resource (UniProt) Knowledgebase and Reference Clusters (http://www.uniprot.org) and the National Center for Biotechnology Information (http://www.ncbi.nlm.nih.gov/).

## Author contributions

M.C., R.P., D.D., F.Z., and A.T.S. designed the research; M.C., R.P., and A.O. performed the research; M.C., R.P., D.D., and A.T.S. analyzed the data; and M.C., R.P., D.D., F.Z., and A.T.S. wrote and edited the paper.

## Additional information

*Supplementary information*. Supplementary information accompanies this paper.

*Competing interests*. The authors declare no competing interests.

## METHODS

### Materials

All materials used for buffer preparation, protein expression, and protein purification were purchased from commercial vendors and were used as received.

### Bioinformatics

All ATE1 sequences were obtained from the Universal Protein Resource (UniProt) Knowledgebase and Reference Clusters (http://www.unprot.org) or the National Center for Biotechnology Information (http://www.ncbi.nlm.nih.gov/). Sequences were retrieved using UniProt ATE1 or arginyl transferase or arginine transferase as an input. Sequences were aligned using EMBL’s online European Bioinformatics Institute (EMBL-EBI) platform containing the Clustal Omega program implementing the ClustalW algorithm ^50,51^ and the Blosum62 matrix.^52^ The sequence output was first visually inspected using the program MEGAX, and the same sequence input was further refined by use of the Multiple Sequence Comparison by Log-Expectation (MUSCLE) algorithm prior to bootstrapping via the Jones-Taylor-Thornton (JTT) method to generate a phylogenetic tree of organismal ATE1 sequences.

### Construct design, protein expression, and protein purification

The cloning of *Mm*ATE1-1 (Uniprot ID Q9Z2A5) followed similar protocols as described in ^23^. The cloning of *Sc*ATE1 (Uniprot ID P16639) followed similar protocols as described in ^18^.

The expression of *Mus musculus* (*Mm*) ATE1-1 followed a similar protocol as described in ^20^. Briefly, *E. coli* BL21 (DE3) codon plus chemically competent cells containing the *Mm*ATE1-1 pET-29a(+) plasmid were grown overnight at 37 °C with shaking at 200 rpm in 100 mL Luria Broth (LB) supplemented with 37 µg/mL (final) chloramphenicol and 50 µg/mL (final) of kanamycin. Next, *ca.* 20 mL of the overnight culture were inoculated into 12 2-L baffled expression flasks pre-charged with 1 L of sterile LB that had been supplemented with 37 µg/mL (final) chloramphenicol and 50 µg/mL (final) of kanamycin. The cells were grown at 37 °C with shaking at 300 rpm until the optical density at 600 nm (OD_600_) reached 0.6-0.8, at which point the flasks were then cold shocked at 4 °C for 2 hr. After cold shocking, isopropyl β-D-1-thiogalactopyranoside (IPTG) was added to a final concentration of 1 mM to each flask to induce protein production; these flasks were then incubated overnight at 18 °C with 200 rpm shaking. After 18-20 hr of incubation, the cells were harvested by centrifuging at 5,000 x*g* for 12 min at 4 °C. Harvested cells were then resuspended in 30 mL resuspension buffer (50 mM Tris pH 7.5, 100 mM NaCl and 5 % (v/v) glycerol). After resuspension the cells were flash frozen in liquid nitrogen and kept at -80 °C.

The purification of *Mm*ATE1-1 followed a similar protocol as described in ^20^. Briefly, frozen cells were first thawed in a water bath and then stirred to homogenize. After the thawed cells were homogenized, solid phenylmethylsulfonylfluoride (PMSF) was added prior to the cells being sonicated by an ultrasonic cell disruptor operating at 4 °C. The lysed cells were spun in an ultracentrifuge at 190,000 x*g* for 1 hour at 4 °C. The supernatant was then applied to a HisTrap HP 5 mL column, washed extensively with 5 CV wash buffer (50 mM Tris pH 8.0, 300 mM NaCl, 10 % (v/v) glycerol, 1 mM TCEP). After several additional washes with wash buffer containing increasing imidazole concentrations*, Mm*ATE1-1 was eluted using 100 % elution buffer (50 mM Tris pH 8.0, 300 mM NaCl, 300 mM imidazole, 10 % (v/v) glycerol). These fractions were concentrated using a Millipore spin concentrator with a molecular weight cutoff (MWCO) of 30 kDa. For tag cleavage, *Mm*ATE1-1 was buffer exchanged into a Tobacco Etch Virus (TEV) cleavage buffer (50 mM Tris pH 7.5, 250 mM KCl, 0.5 mM EDTA, 5% (v/v) glycerol, 1 mM TCEP). To the sample, homemade TEV protease was added in a ratio of 40 µg of protease per 1 mg of *Mm*ATE1-1 protein, and the mixture was rocked overnight at 4°C. The mixture of protease and cleaved protein was then concentrated, and the sample was applied to a Superdex 200 size-exclusion gel-filtration column pre-equilibrated with SEC buffer (50mM Tris pH 7.5, 250mM KCl, 5%v/v glycerol, 1mM DTT). For uncleaved protein, the HisTrap HP-purified and concentrated *Mm*ATE1-1 protein was applied directly to the gel-filtration column. In both cases (cleaved or uncleaved), protein corresponding to a monomeric *Mm*ATE1-1 was pooled, concentrated, quantified using the DC Lowry assay, and characterized using 15% SDS-PAGE analysis. For the cleaved protein, loss of the (His)_6_ tag was verified via Western blotting using a commercially available anti-(His)_6_ antibody (MilliporeSigma).

The purification of *Sc*ATE1 followed a similar protocol as described in^18^, which was nearly identical to that of *Mm*ATE1-1 with the following modifications: 1) the gene encoding for *Sc*ATE1 was subcloned into the pET-21a(+) plasmid; 2) *Sc*ATE1 was electroporated and expressed in *E. coli* BL21(DE3) electrocompetent cells; 3) colony selection, starter cultures, and large-scale expression were all accomplished with transformed cells grown in the presence of 100 µg/mL of ampicillin (final concentration); 4) the (His)_6_ tag was not cleaved from *Sc*ATE1 prior to additional analyses.

### AlphaFold Modeling

All AlphaFold models were obtained from the predicted AlphaFold Protein Structure Database ^53^ or modeled *de novo* using the AlphaFold3 structural prediction server.^54^ The following AlphaFold model IDs or UniProt IDs were used to obtain the predicted structures as well as the predicted, aligned error: UniProt IDs: Q9Z2A5-1, Q9Z2A5-2, Q9Z2A5-3, Q9Z2A5-4 (*Mm*ATE1 isoforms -1 through -4, respectively), and AlphaFold Protein Structure Database IDs: J3QNU1 (*Mm*ATE1-4), Q80YP1 (*Mm*ATE1-2), P16639 (*Sc*ATE1).

### Size-exclusion coupled small-angle X-ray scattering (SEC-SAXS)

A 10x buffer (500 mM Tris, pH 7.5, 2.50 M KCl, 20 % (v/v) glycerol, 10 mM DTT) was first prepared for each SEC-SAXS run. For all analyses, a purified protein sample (*e.g.*, uncleaved *Mm*ATE1-1, cleaved *Mm*ATE1-1, uncleaved *Sc*ATE1) was re-polished by running prior prepared protein along an analytical Superdex200 column in 1x buffer from the 10x stock. Parsimonious samples of each protein were then collected, the concentrations were measured using a DC Lowry assay, and SDS-PAGE was run to assure a high sample purity. Protein samples were then diluted to a range of 2 - 8 mg/mL in 1 mg/mL increments before the samples were flash frozen and shipped on dry ice. At the SIBLYS beamline (ALS beamline 12.3.1), samples were screened after passage along a PROTEIN KW-803 column equilibrated with SAXS buffer using an autosampler. Eluent was split 2:1 between the X-ray synchrotron radiation source (SAXS) and a series of four inline analytical instruments: 1) Agilent 1260 series multiple wavelength detector (MWD); 2) Wyatt Dawn Helos multi-angle light scattering detector; 3) Wyatt DynaPro Titan quasi-elastic light scattering detector; and 4) Wyatt Optilab rEX refractometer. An incident light (λ = 1.03 Å) with a set sample-to-detector distance of 1.5 m led to scattering vectors, q, these vectors ranged from 0.01 Å^−1^ to 0.5 Å^−1^ the vector q is defined as q = 4πsinθ/λ where 2θ is the measured scattering angle. Data were collected during 40 min with 3 s exposures. SEC-SAXS chromatograms were generated and initial SAXS curves were analyzed using RAW, with arbitrary data scaling. UV, MALS, QELS, and differential refractive index data were also collected and analyzed. The program RAW was used to generate Guinier and Kratky plots. ^55,56^ Values including molecular weight, radius of gyration (*R_g_*) and the maximum particle dimension (*D_max_*) were determined. *Ab initio* molecular envelopes were generated using the ATSAS package.^57^ DAMMIF/N results were displayed using UCSF ChimeraX.^58^ To analyze flexibility within the *Mm*ATE1-1 IDR region, BilboMD classic,^59^ which uses the Chemistry at HARvard Macromolecular Mechanics (CHARMM) simulation program, was used to generate molecular dynamic models. The AlphaFold *Mm*ATE1-1 model (AF-Q9Z2A5-F1) PDB file, the predicted aligned error (pae) file, and the experimental three-column SAXS .dat file were all used as imputs. The BilboMD ouput generated 11 predicted models, which were averaged to generate theoretical scattering data that were then compared to the experimental scattering data through the FoXS server.

### Sample Preparation for hydrogen deuterium exchange (HDX)

Size-exclusion purified samples (*Mm*ATE1-1 both cleaved and uncleaved or *Sc*ATE1 uncleaved) were thawed and diluted to 10 μM concentration. A deuterated buffer was made of 50 mM Tris in D_2_O at pD 7.5, 250 mM KCl, 5 % (v/v) glycerol, and 1 mM DTT. A quench solution was made containing 100 mM glycine, 0.8 M urea and 20 mM TCEP at pH 2.5. Deuterium exchange was initiated by mixing 2 µL of 10 µM protein with 38 µL of D_2_O buffer, and deuterium exchanged was quenched with 60 µL of quench solution after 10, 100, and 1000 s deuterium exchange, respectively, and the quenched samples were then placed on ice. Fully labeled control samples were prepared by labeling ATE1 in deuterated buffer after denaturation of *Mm*ATE1-1 (6 M deuterated urea, pH 2.3) for 3 hr at 25°C, or after denaturation of *Sc*ATE1 (6 M deuterated GndHCL, pH 2.3) for 24 hr at 4 °C. Non-deuterated controls were also run using a non-deuterated buffer and treated the same as all deuterated samples. All controls and sample time points were repeated in triplicate and analyzed for consistent behavior.

### HDX-Mass Spectrometry (HDX-MS) measurements and data anlysis

In preparation for mass spectrometry (MS) analyses, samples were incubated on ice for 1 min after each quench before each sample was injected into a refrigerated (0 °C) Waters HDX nanoACQUITY UPLC (Waters, Milford, MA) coupled to a hybrid Q-ToF Synapt G2S mass spectrometer. Online digestion was completed at 20 °C with a constant sample flow rate of 200 µL/min in a separate chamber using an IDEX column packed in-house with immobilized pepsin on agarose resin beads (Thermo Scientific, Pierce). The generated peptides were trapped with an Acquity UPLC BEH C18 peptide 1.7 μm VanGuard column (Waters) and desalted for 3 min with solvent A (0.1 % (v/v) formic acid in Milli-Q water, pH 2.5). The samples were run for 7 min with a linear 40 μL/min flow rate with increasing solvent B (acetonitrile with 0.1 % (v/v) formic acid) across a gradient of 5% to 35 % (v/v) solvent B. Peptides were eluted into a Synapt G2S mass spectrometer with an electrospray ionization source, which was operated in a positive ionization mode with a capillary voltage of 3 kV, a source temperature of 80 °C, and a sampling cone of 30 kV. For lock-mass calibration, human Leu-enK (Sigma-Aldrich) was recorded throughout the analysis to ensure mass accuracy.

For peptide identification, nondeuterated samples of ATE1 were analyzed using the same UPLC method and peptides were fragmented by collision-induced dissociation (CID) using MSe acquisition mode. To identify ATE1 peptides, data were processed using the ProteinLynx Global Server (PLGS) version 3.0.3 (Waters), with the sequence of *Mm*ATE1-1 or *Sc*ATE1. To filter peptides identified, a minimum peptide score of 7 and a minimum of 0.3 fragmentation products per amino acid were utilized as filtration parameters. Raw MS data were imported into DynamX 3.0, and all identified peptides were manually checked in DynamX version 3.0. (Waters); mass spectra with a signal-to-noise ratio (S/B) less than 10 were excluded from all analyses. For *Sc*ATE1, HD Examiner version 3.3 was used to analyze FD peptides.

The percent deuterium uptake of peptides presented in this work is the average of at least three technical replicates. The three deuteration time points were acquired in triplicate. Fully deuterated controls were acquired for normalization purposes. The normalized percent deuterium uptake (%D) for each peptide, at incubation time *t*, was calculated as described in the equation in the following:

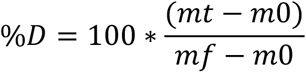

where mt, m0, and mf are the centroid masses at incubation time *t*, the undeuterated control, and the fully deuterated control, respectively. For visualization, HDX results were mapped onto the *Mm*ATE1-1 AlphaFold model generated in this work, the *Sc*ATE1 high-resolution X-ray crystal structure (PDB ID 7TIF), and/or the *Sc*ATE1 AlphaFold model (AF-P16639-F1 in order to visualize the putative positions of the regions otherwise disordered in the X-ray structure) using ChimeraX.^58^ The HDX uptake plots are included in the Supporting Information.

## Data availability

Data are available from the corresponding author upon request.

## Abbreviations

ATE1: arginyltransferase 1
GNAT: GCN5-related *N*-acetyltransferase fold
HDX-MS: hydrogen deuterium exchange mass spectrometry
IDR: intrinsically disordered region
JTT: Jones-Taylor-Thornton
MALS: multi-angle light scattering
*Mm*: *Mus musculus*
MSA: multiple sequence alignment
MUSCLE: multiple sequence comparison by log-expectation
PTM: post-translational modification
SAXS: small angle X-ray scattering
*Sc*: *Saccharomyces cerevisiae*
SEC: size-exclusion chromatography
tRNA: transfer ribonucleic acid

