## Supporting Information for "Identification of an intrinsically disordered region (IDR) in arginyltransferase 1 (ATE1)"

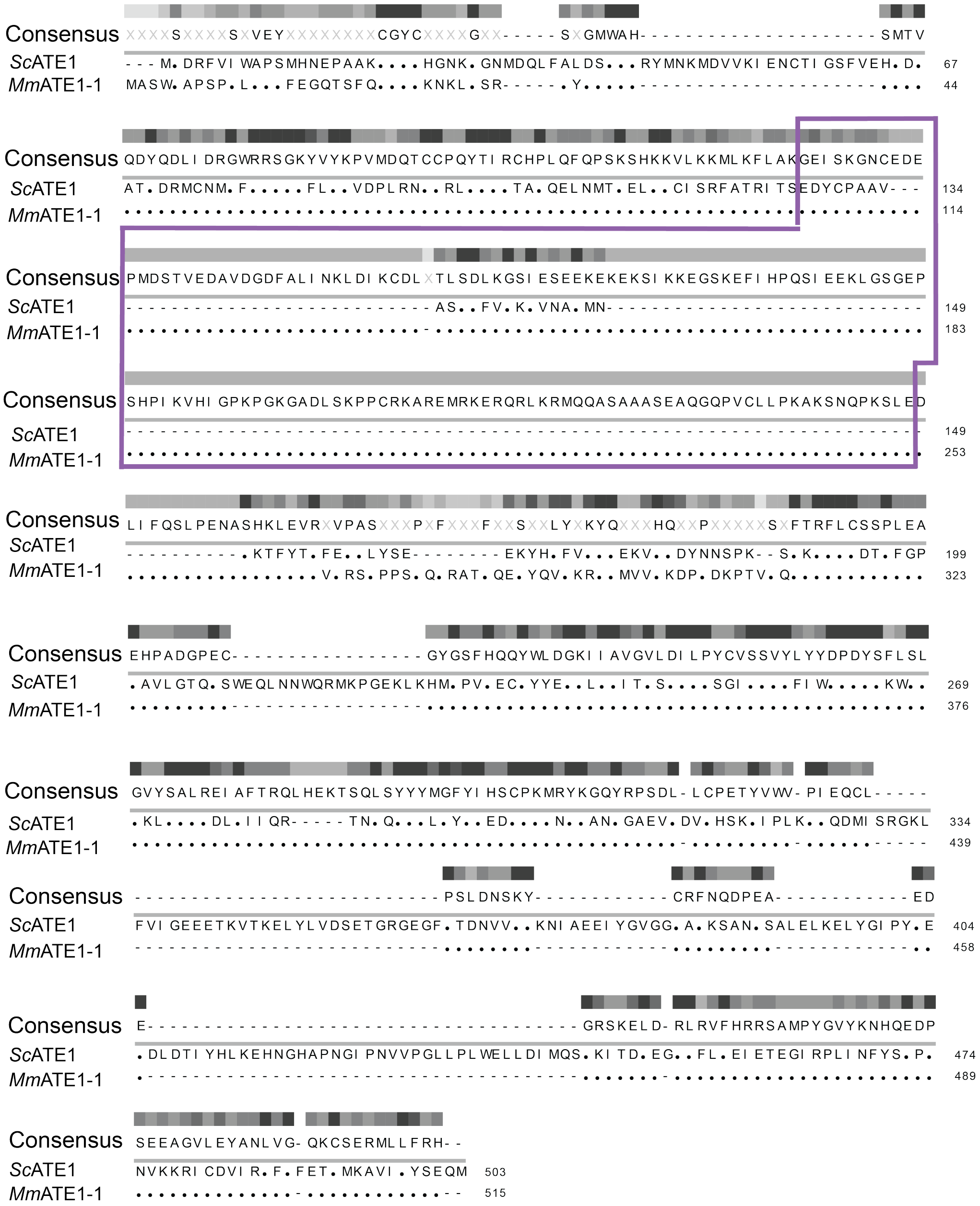

**Figure S1**. Sequence alignment of *Mm*ATE1-1 and *Sc*ATE1. The region corresponding to the IDR in *Mm*ATE1-1 is highlighted with a purple box and is missing in *Sc*ATE1, as evidenced by the dashed lines.

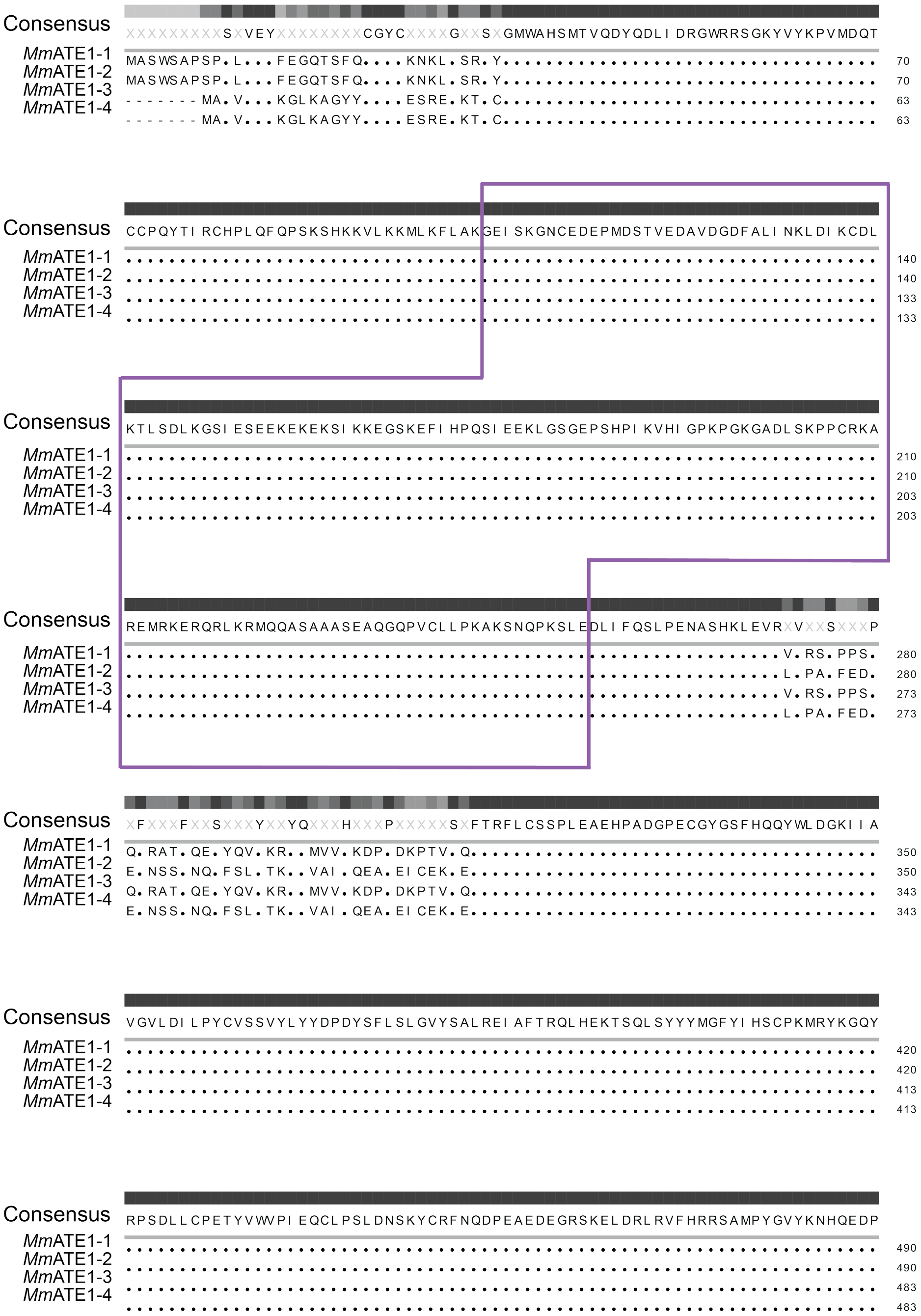

**Figure S2**. Multiple sequence alignment (MSA) of *Mm*ATE1-1 through -4 isoforms. The region highlighted with the purple box represents the IDR, which is 100% conserved amongst the four major isoforms.

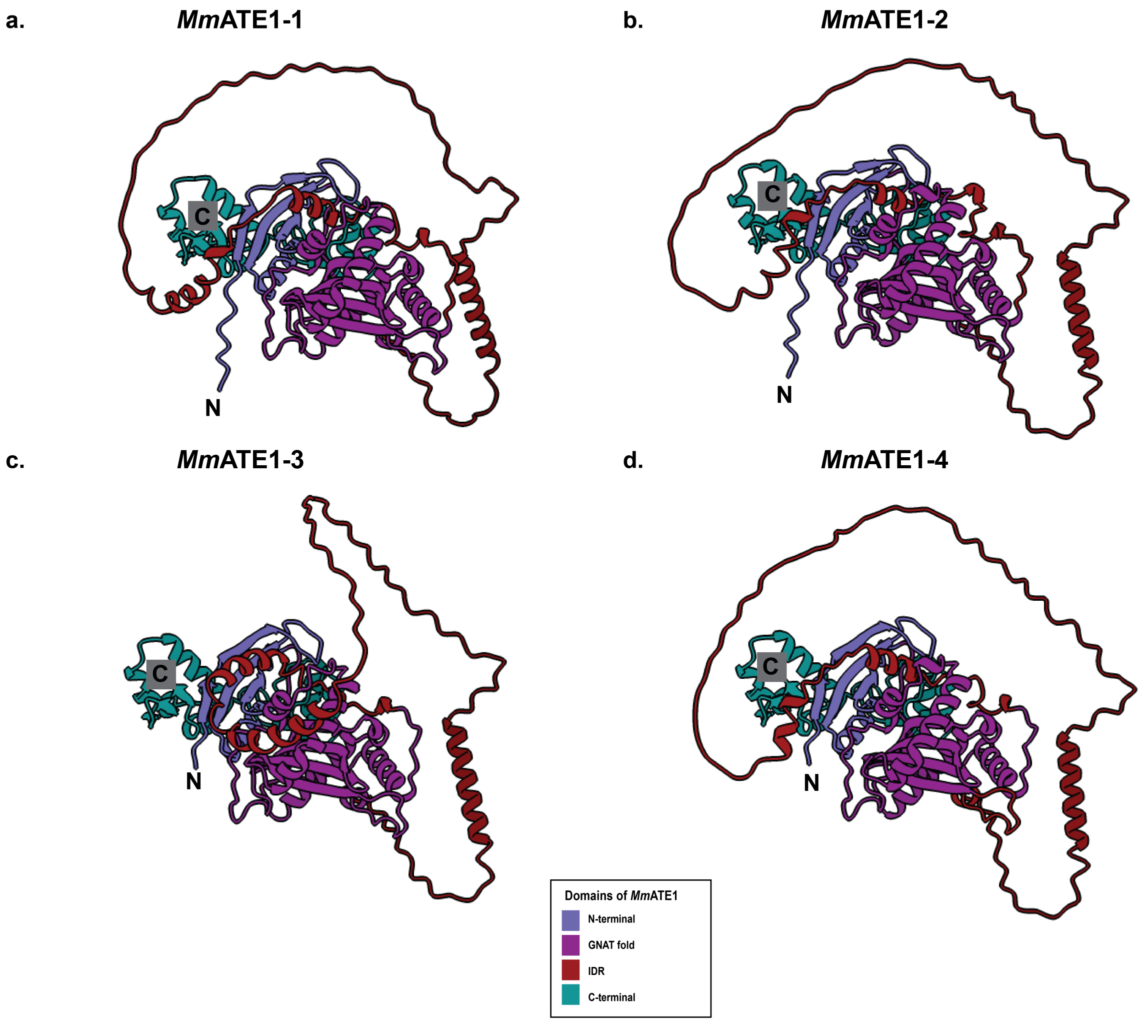


**Figure S3**. Lowest-energy calculated AlphaFold3 models of *Mm*ATE1 isoforms 1 (**a**), 2 (**b**), 3 (**c**), and 4 (**d**) illustrating conservation of the IDR as well as other key ATE1 domains. Due to its dynamic nature, the IDR (colored in brick red) appears in slightly different conformations among the predicted structures of the different isoforms. The N-terminal domains of *Mm*ATE1-3 and *Mm*ATE1-4 (colored in royal blue) are shorter due to a shorter sequence in exon 1 in these isoforms (see Fig. S2). The labels 'N' and 'C' represent the locations of the N- and C-termini, respectively.

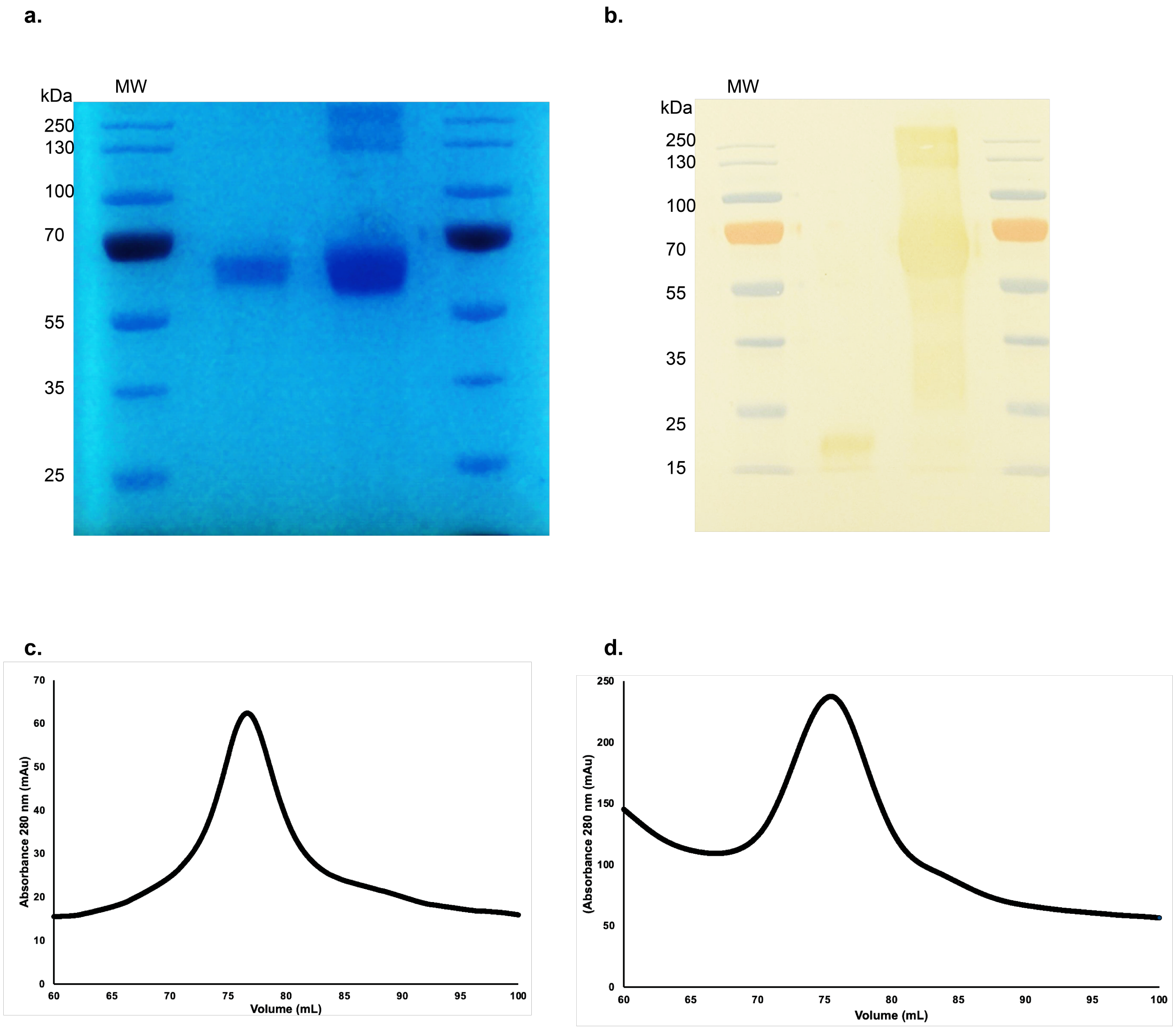


**Figure S4**. Analysis of purified *Mm*ATE1-1. **a.** SDS-PAGE analysis of purified cleaved (left lane) and uncleaved (right lane) *Mm*ATE1-1 after size-exclusion chromatography (SEC). **b**. Western blot analyses of the samples in panel **a**. using an anti-$(His)_6$ antibody demonstrating the cleavage of *Mm*ATE1-1 in the left lane. Size-exclusion chromatograms of purified, cleaved (**c**) and uncleaved (**d**) *Mm*ATE1-1. In both cases, the primary peak, centered at *ca*. 77 mL elution volume corresponds to monomeric *Mm*ATE1-1 based on calibration standards.

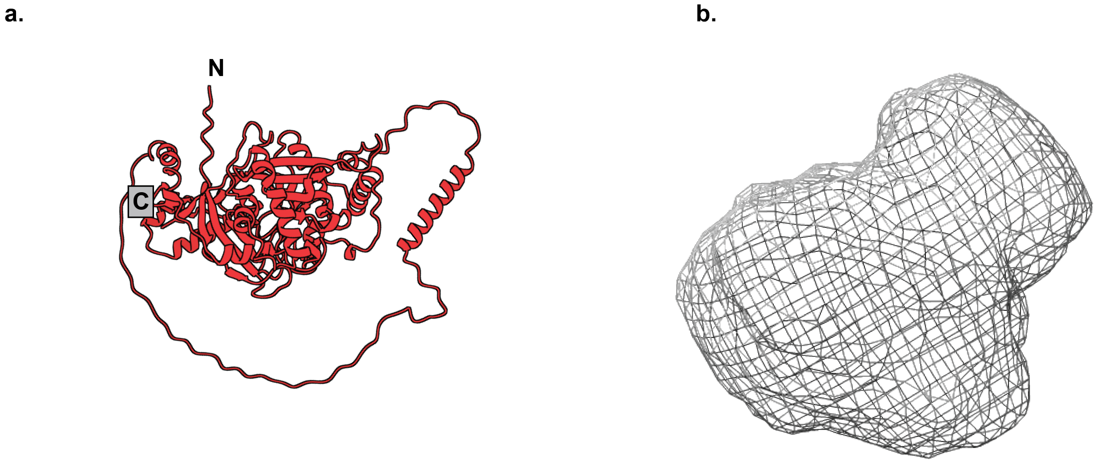


**Figure S5**. Comparison of the lowest-energy *Mm*ATE1-1 AlphaFold3 model (**a**) to that of the *ab initio* envelope of cleaved *Mm*ATE1-1 generated from the SEC-SAXS data (**b**). The labels 'N' and 'C' represent the locations of the N- and C-termini, respectively.

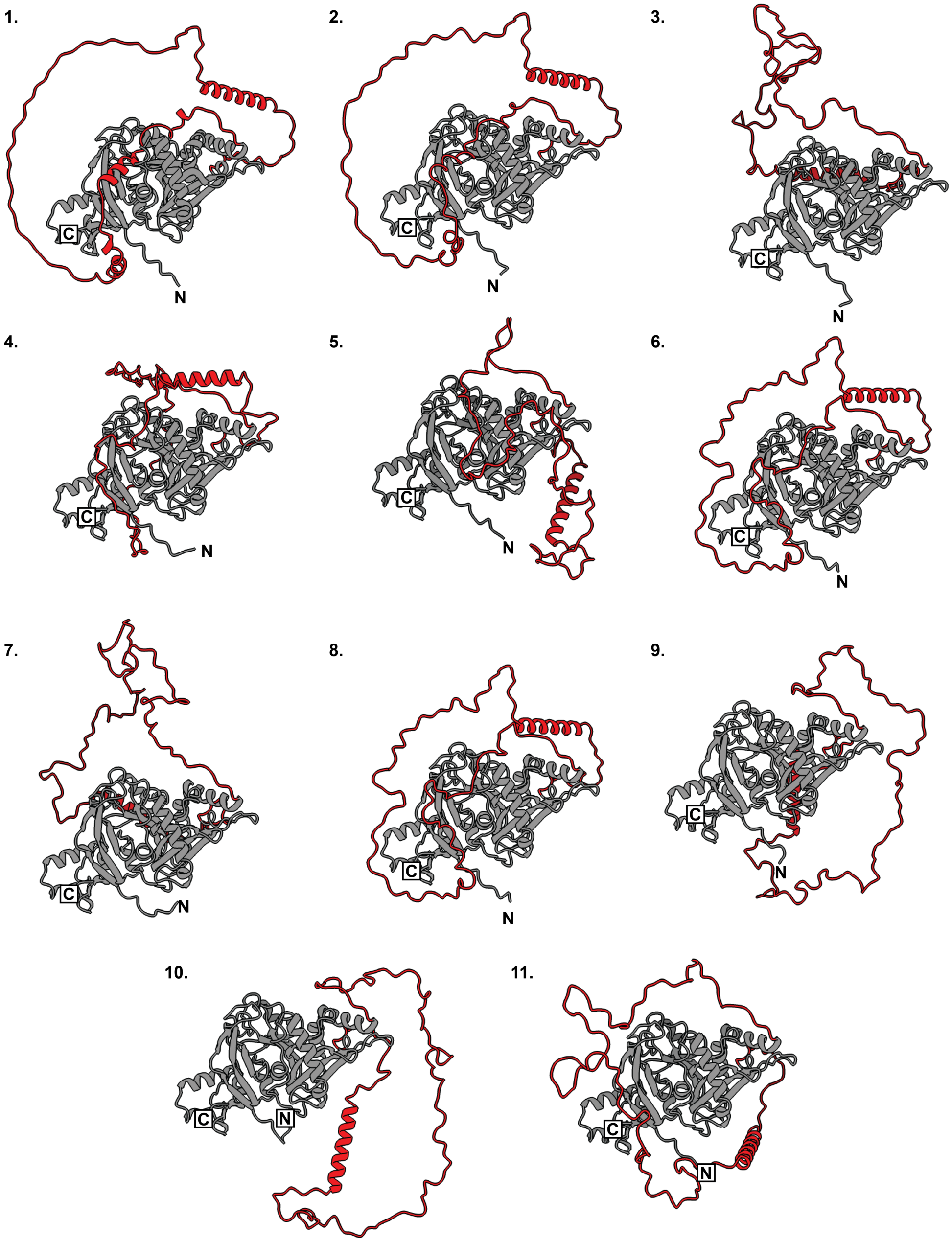


**Figure S6**. BilboMD analysis of the *Mm*ATE1-1 SAXS data coupled with the *Mm*ATE1-1 AlphaFold model illustrates flexibility in the IDR. Eleven (11) different conformations were sampled to recapitulate the experimental SAXS data, and major conformational changes in the IDR (colored in red) occurred during the simulations. The BilboMD analysis also predicted movement of the N-terminal region, but these changes are far less drastic than the IDR. The labels 'N' and 'C' represent the locations of the N- and C-termini, respectively.

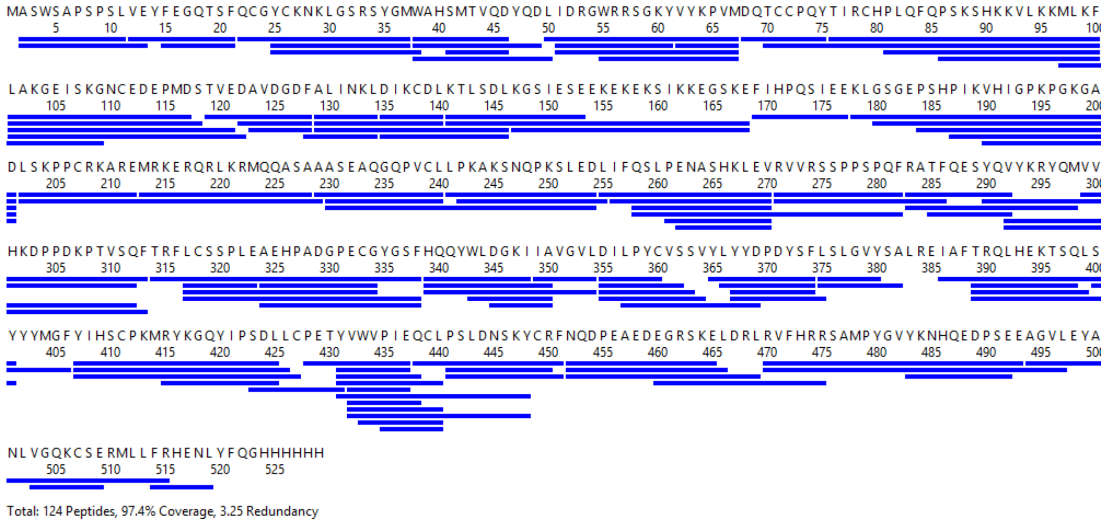

**Figure S7**. Sequence coverage map of *Mm*ATE1-1. Solid blue lines denote the peptic fragments analyzed in this study with total sequence coverage of 97.4%

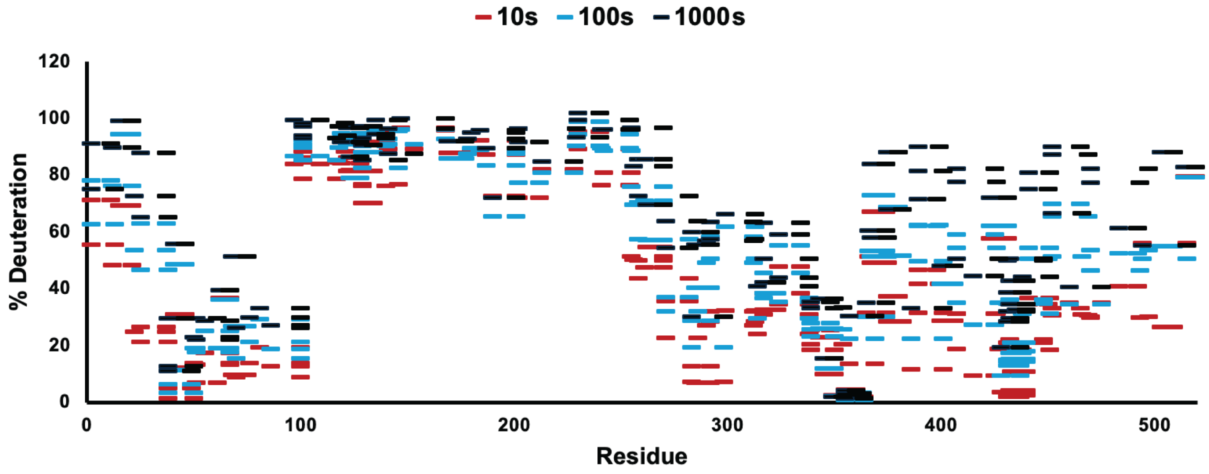


**Figure S8**. Woods plot of *Mm*ATE1-1 representing percent deuteration values in the apo state for time points 10s (red), 100s (blue), and 1000s (black), plotted against amino acid sequence (bottom).

**MMATE1 2-11: ASWSAPSPSL (#1)**

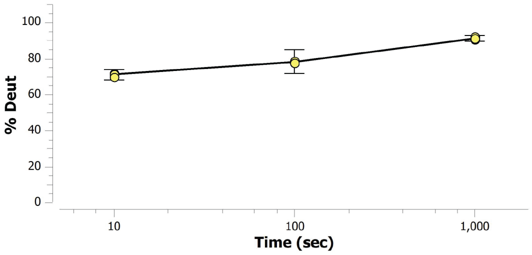


**MMATE1 2-13: ASWSAPSPSLVE (#2)**

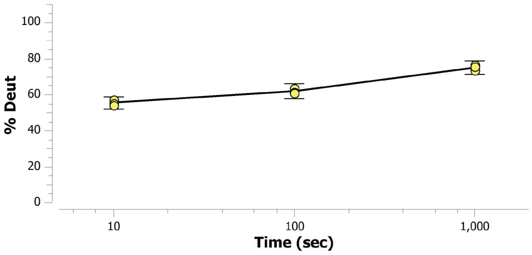


**MMATE1 12-21: VEYFEGQTSF (#3)**

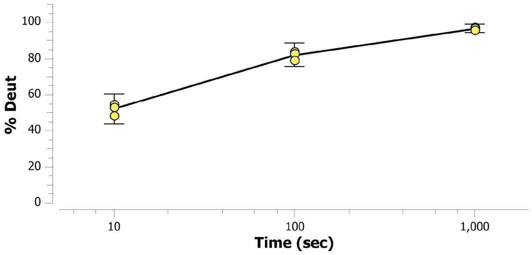


**MMATE1 15-21: FEGQTSF (#4)**

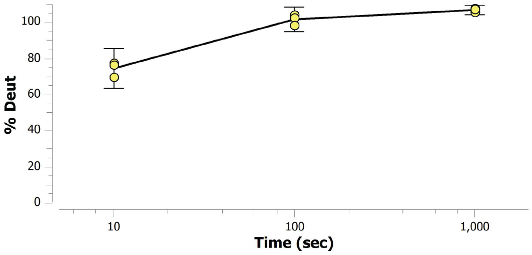


**MMATE1 22-37: QCGYCKNKLGSRSYGM (#5)**

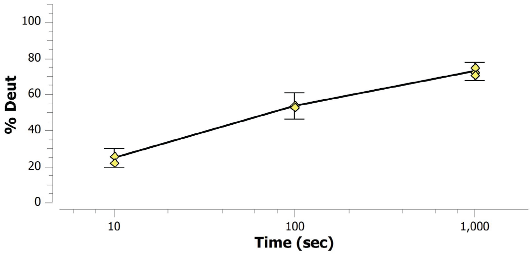


**MMATE1 25-37: YCKNKLGSRSYGM (#6)**

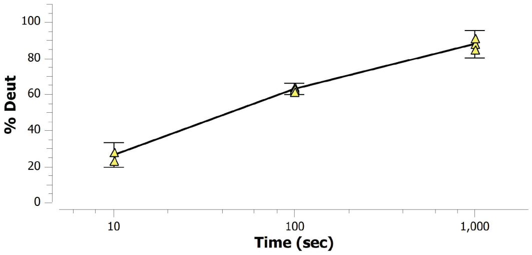


**MMATE1 25-38: YCKNKLGSRSYGMW (#7)**

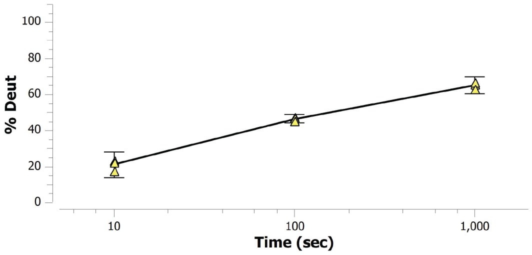


**MMATE1 38-46: WAHSMTVQD (#8)**

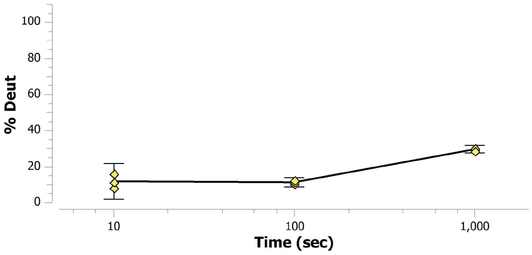


**MMATE1 38-49: WAHSMTVQDYQD (#9)**

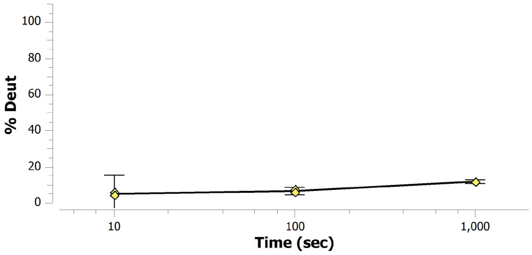


**MMATE1 38-50: WAHSMTVQDYQDL (#10)**

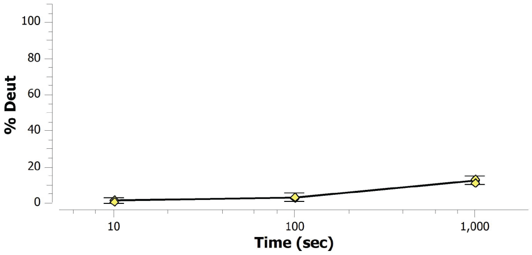

MMATE1 41-46: SMTVQD (#11)

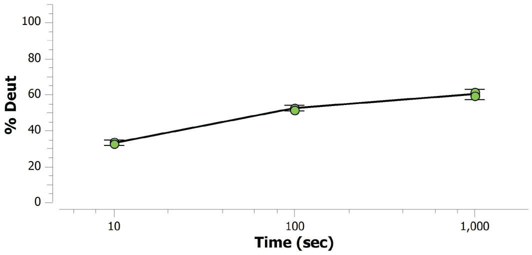


MMATE1 50-67: LIDRGWRRSGKYVYKPVM (#12)

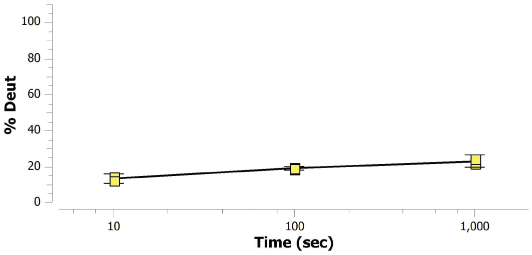


MMATE1 51-61: IDRGWRRSGKY (#13)

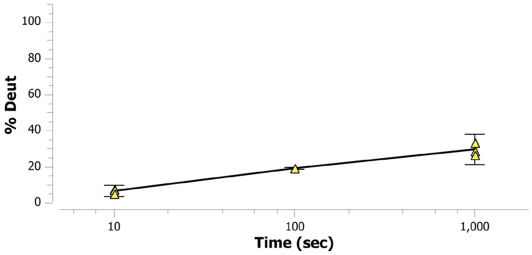


MMATE1 51-67: IDRGWRRSGKYVYKPVM (#14)

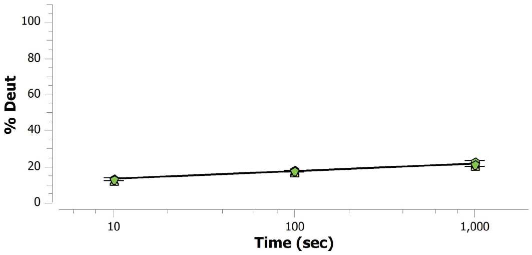


MMATE1 55-67: WRRSGKYVYKPVM (#15)

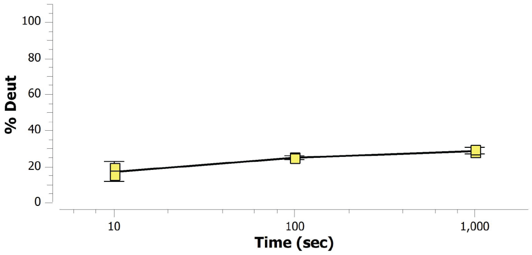


MMATE1 62-67: VYKPVM (#16)

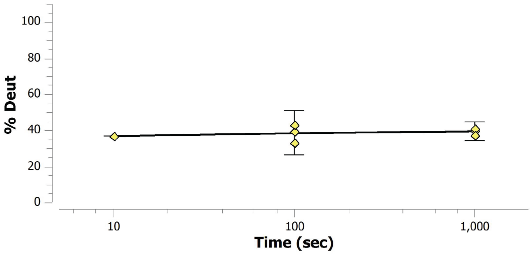


MMATE1 68-75: DQTCCPQY (#17)

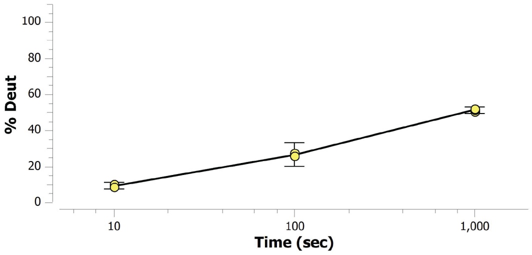


MMATE1 70-100: TCCPQYTIRCHPLQFQPSKSHKKVLKKMLKF (#18)

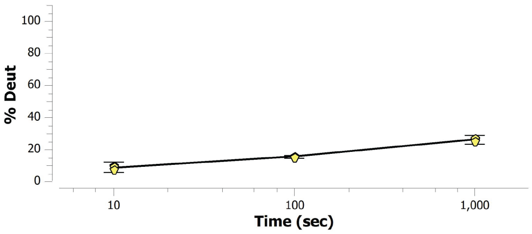


MMATE1 76-100: TIRCHPLQFQPSKSHKKVLKKMLKF (#19)

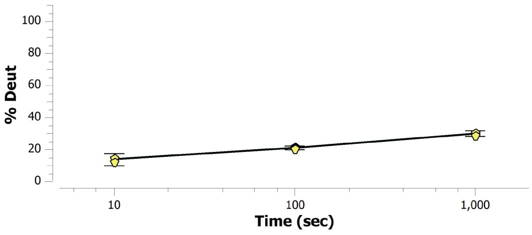


MMATE1 81-100: PLQFQPSKSHKKVLKKMLKF (#20)

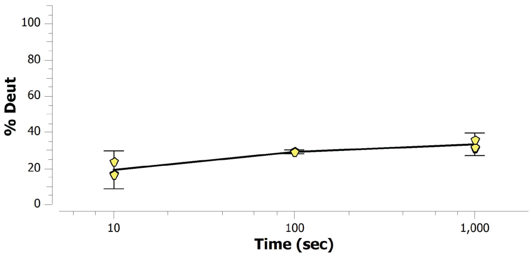

**MMATE1 86-100: PSKSHKKVLKKMLKF (#21)**

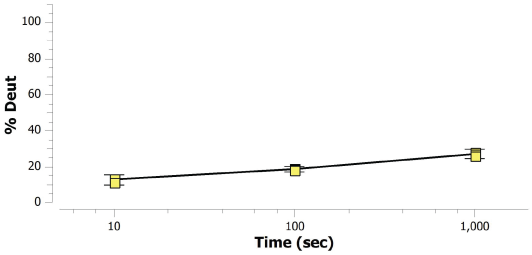


**MMATE1 97-109: MLKFLAKGEISKG (#22)**

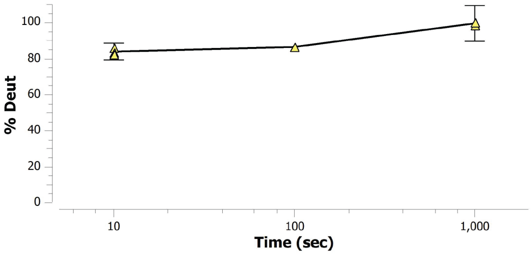


**MMATE1 101-117: LAKGEISKGNCEDEPMD (#23)**

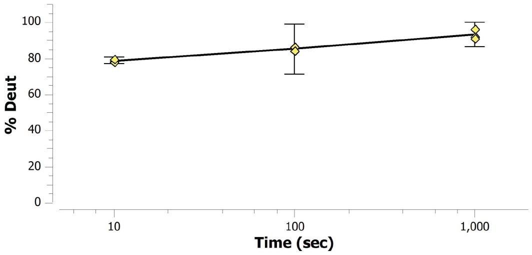


**MMATE1 101-118: LAKGEISKGNCEDEPMDS (#24)**

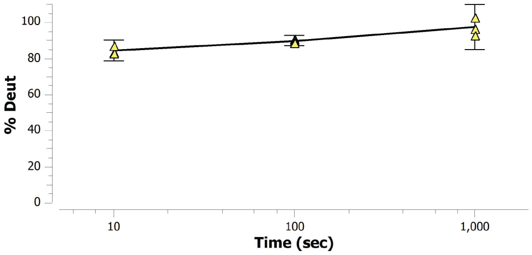


**MMATE1 101-121: LAKGEISKGNCEDEPMDSTVE (#25)**

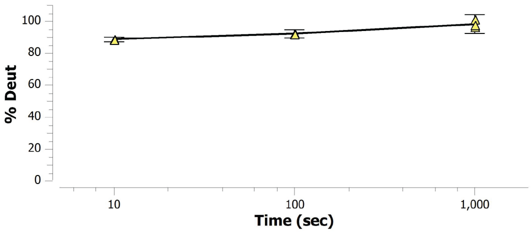


**MMATE1 101-122: LAKGEISKGNCEDEPMDSTVED (#26)**

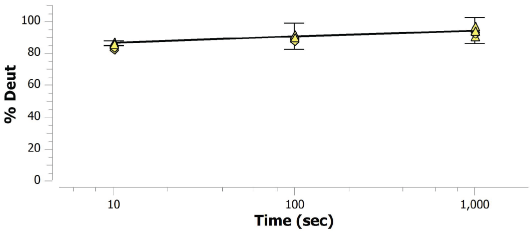


**MMATE1 119-128: TVEDAVDGDF (#27)**

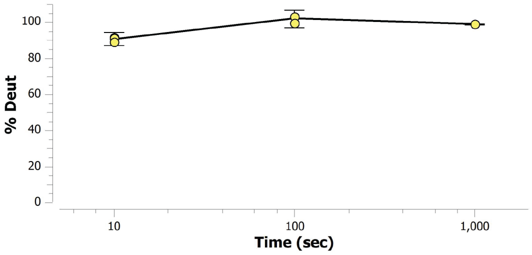


**MMATE1 122-128: DAVDGDF (#28)**

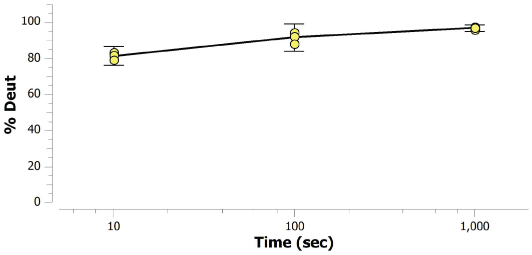


**MMATE1 123-128: AVDGDF (#29)**

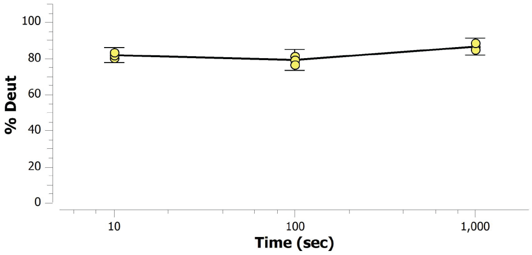


**MMATE1 128-134: FALINKL (#30)**

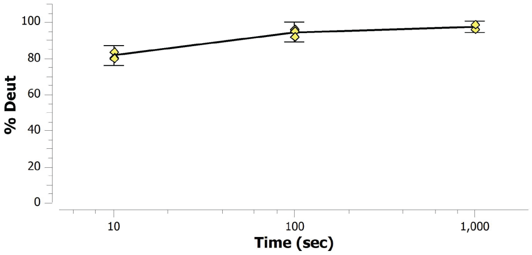

**MMATE1 129-134: ALINKL (#31)**

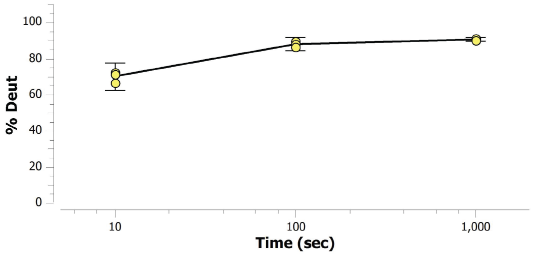


**MMATE1 129-140: ALINKLDIKCDL (#32)**

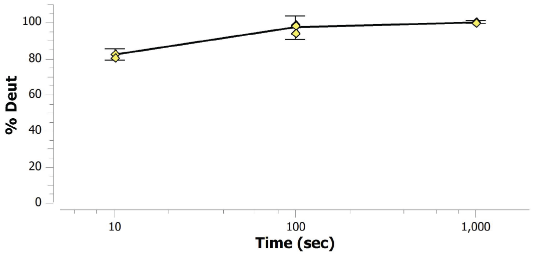


**MMATE1 129-146: ALINKLDIKCDLKTLSDL (#33)**

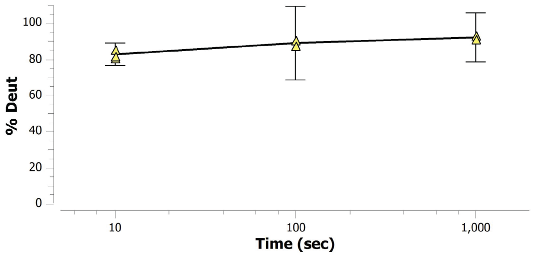


**MMATE1 135-140: DIKCDL (#34)**

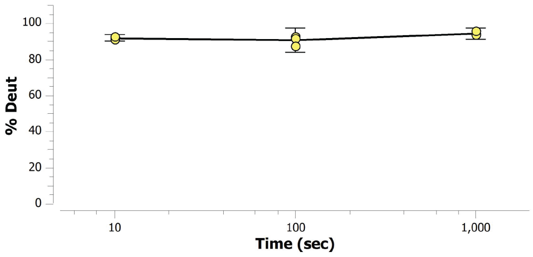


**MMATE1 135-146: DIKCDLKTLSDL (#35)**

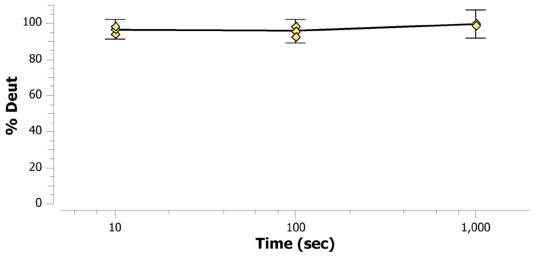


**MMATE1 141-153: KTLSDLKGSIESE (#36)**

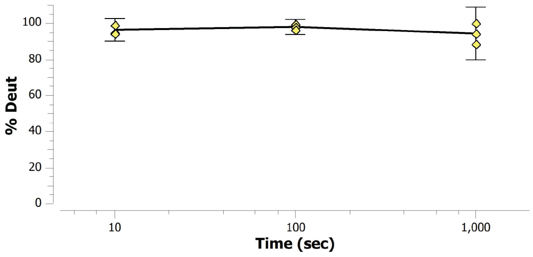


**MMATE1 141-168: KTLSDLKGSIESEEKEKEKSIKKEGSKE (#37)**

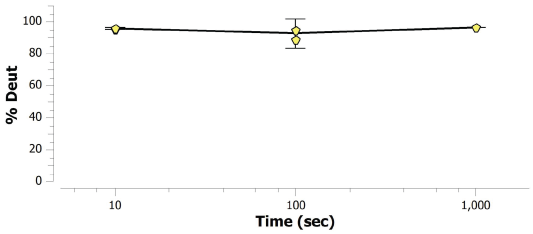


**MMATE1 147-168: KGSIESEEKEKEKSIKKEGSKE (#38)**

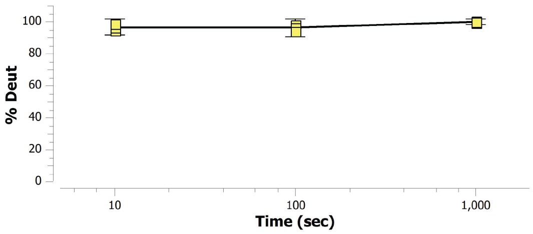


**MMATE1 169-177: FIHPQSIEE (#39)**

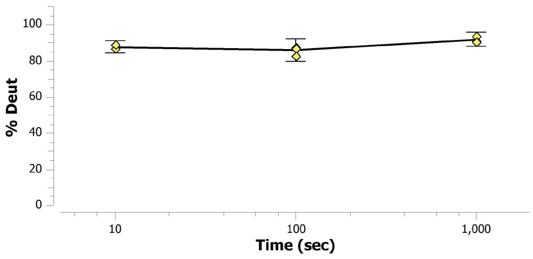


**MMATE1 178-201: KLGSGEPSHPIKVHIGPKPGKGAD (#40)**

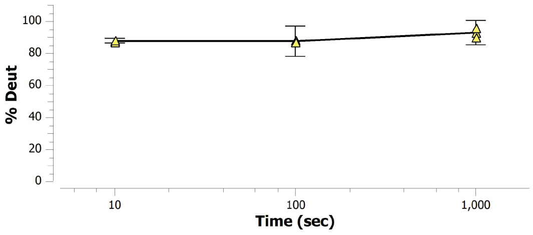

**MMATE1 180-201: GSGEPSHPIKVHIGPKPGKGAD (#41)**

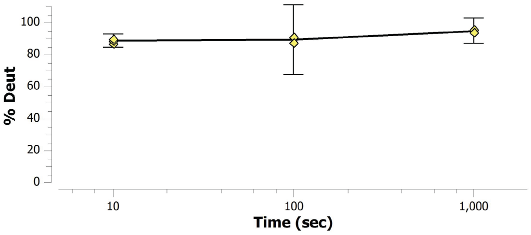


**MMATE1 184-201: PSHPIKVHIGPKPGKGAD (#42)**

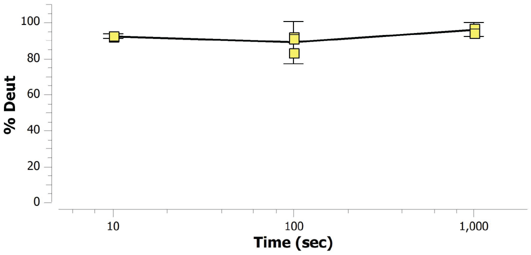


**MMATE1 187-201: PIKVHIGPKPGKGAD (#43)**

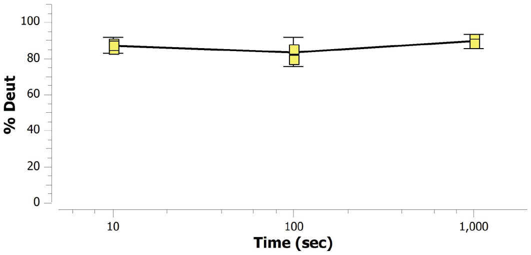


**MMATE1 190-201: VHIGPKPGKGAD (#44)**

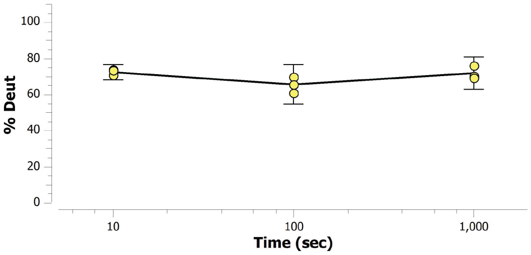


**MMATE1 202-212: LSKPPCRKARE (#45)**

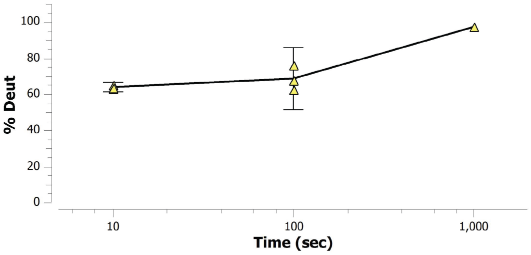


**MMATE1 202-229: LSKPPCRKAREMRKERQRLKRMQQASAA (#46)**

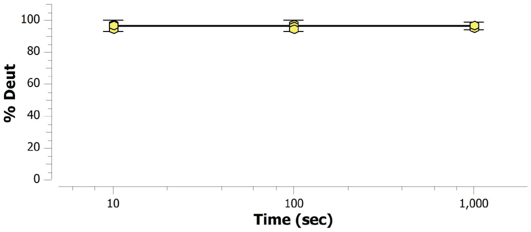


**MMATE1 213-228: MRKERQRLKRMQQASA (#47)**

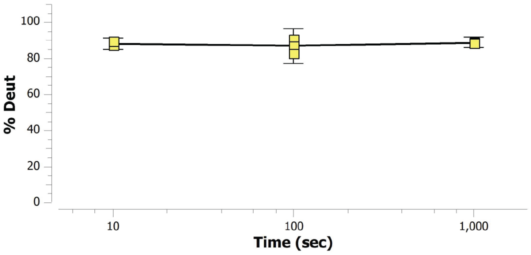


**MMATE1 229-240: AASEAQGQPVCL (#48)**

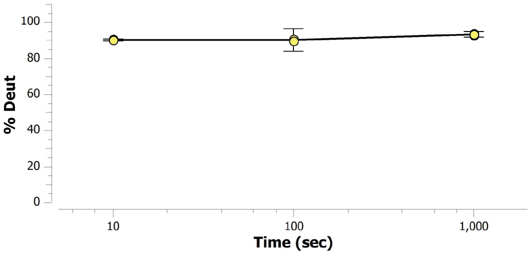


**MMATE1 230-240: ASEAQGQPVCL (#49)**

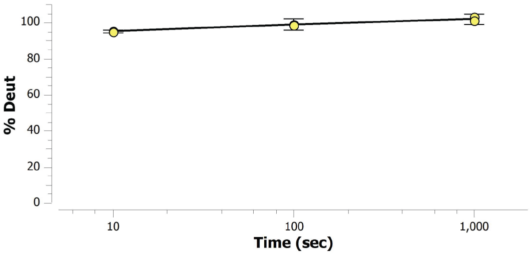


**MMATE1 230-254: ASEAQGQPVCLLPKAKSNQPKSLED (#50)**

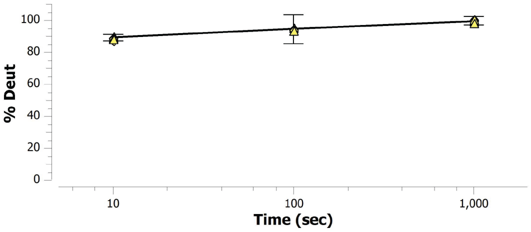

**MMATE1 241-254: LPKAKSNQPKSLED (#51)**

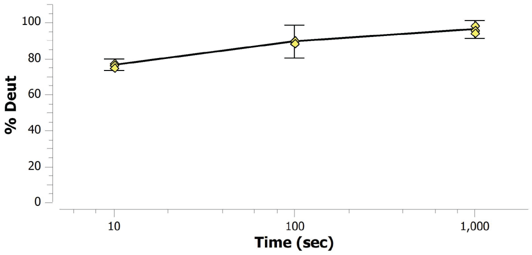


**MMATE1 242-255: PKAKSNQPKSLEDL (#52)**

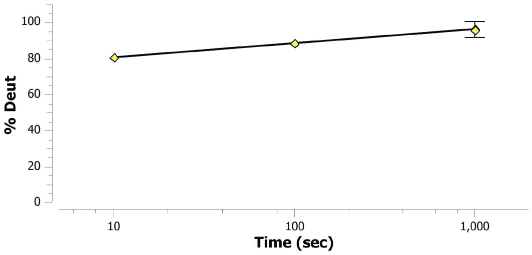


**MMATE1 255-270: LIFQSLPENASHKLEV (#53)**

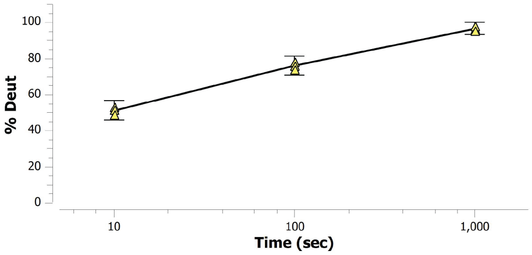


**MMATE1 256-270: IFQSLPENASHKLEV (#54)**

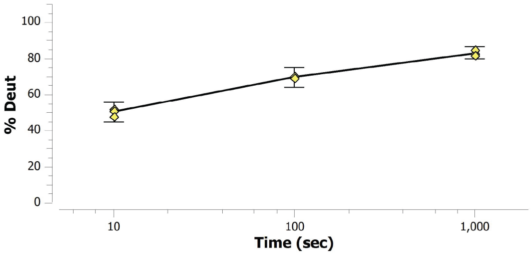


**MMATE1 258-270: QSLPENASHKLEV (#55)**

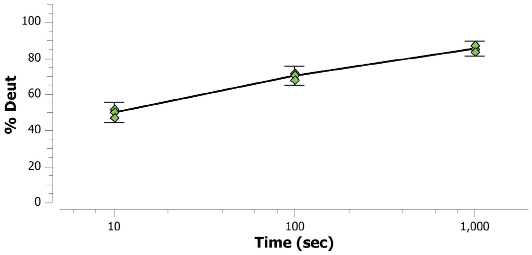


**MMATE1 258-282: QSLPENASHKLEVRVVRSSPPSPQF (#56)**

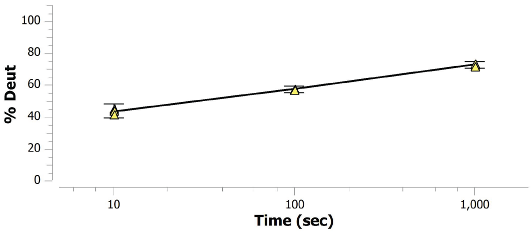


**MMATE1 261-270: PENASHKLEV (#57)**

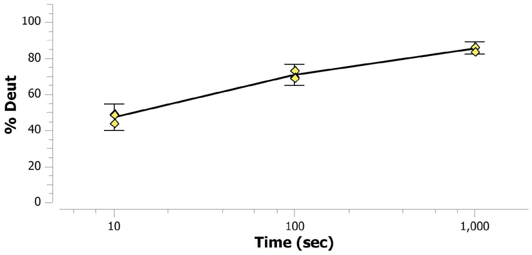


**MMATE1 262-270: ENASHKLEV (#58)**

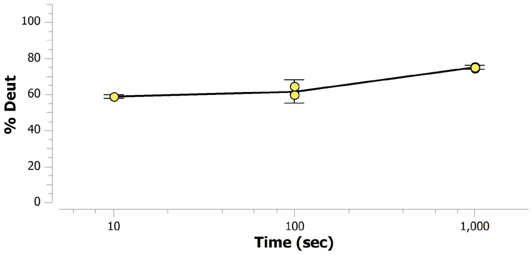


**MMATE1 271-282: RVVRSSPPSPQF (#59)**

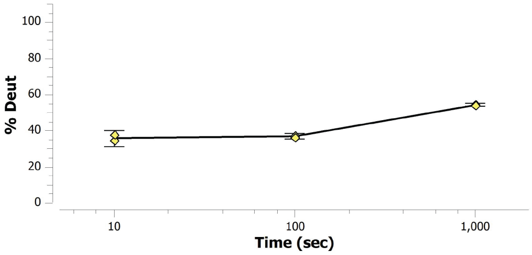


**MMATE1 271-286: RVVRSSPPSPQFRATF (#60)**

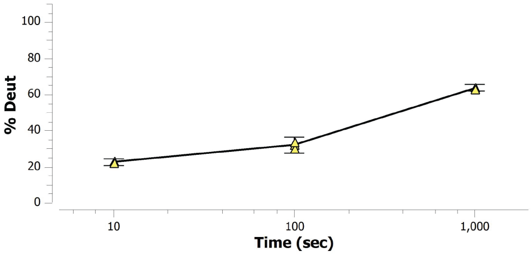

**MMATE1 283-292: RATFQESYQV (#61)**

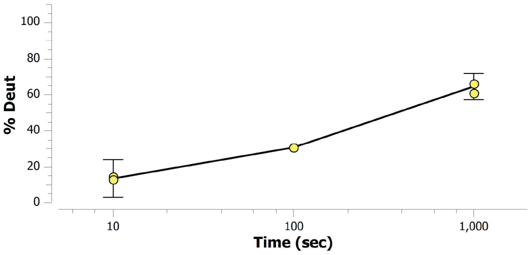


**MMATE1 283-298: RATFQESYQVYKRYQM (#62)**

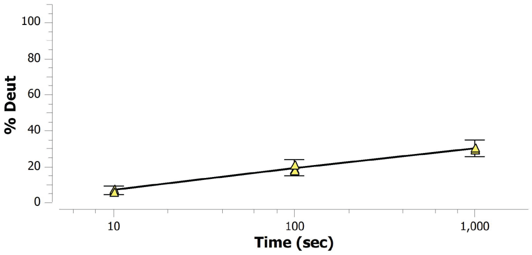


**MMATE1 285-292: TFQESYQV (#63)**

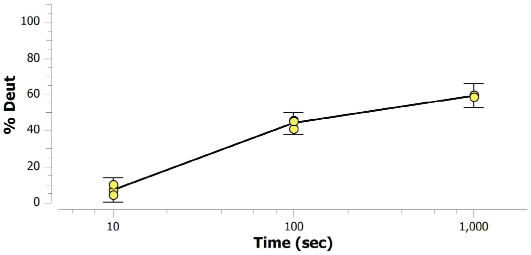


**MMATE1 290-312: YQVYKRYQMVVHKDPPDKPTVSQ (#64)**

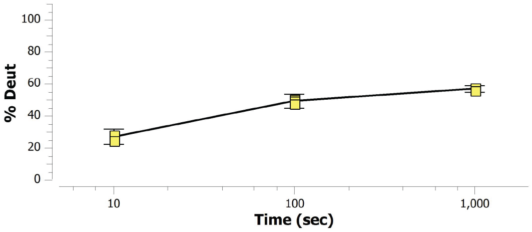


**MMATE1 292-312: VYKRYQMVVHKDPPDKPTVSQ (#65)**

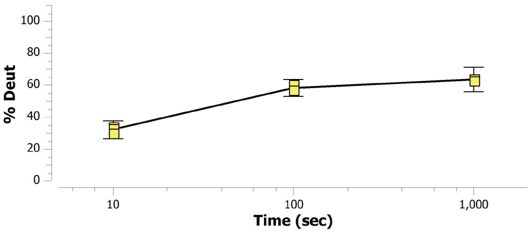


**MMATE1 292-313: VYKRYQMVVHKDPPDKPTVSQF (#66)**

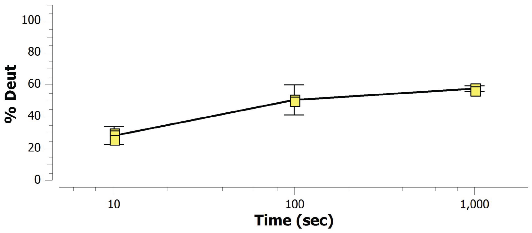


**MMATE1 299-313: VVHKDPPDKPTVSQF (#67)**

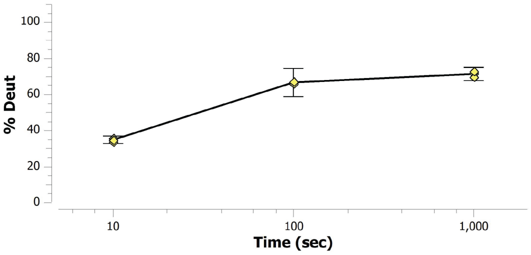


**MMATE1 314-338: TRFLCSSPLEAEHPADGPECGYGSF (#68)**

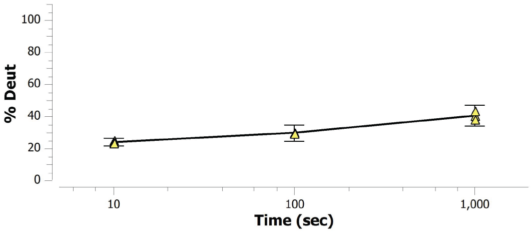


**MMATE1 317-323: LCSSPLE (#69)**

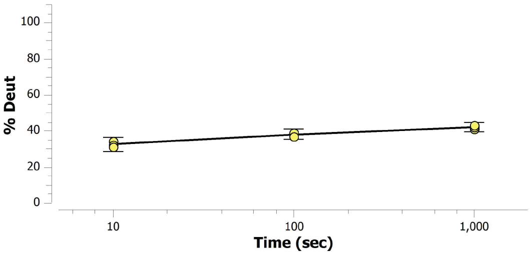


**MMATE1 317-334: LCSSPLEAEHPADGPECG (#70)**

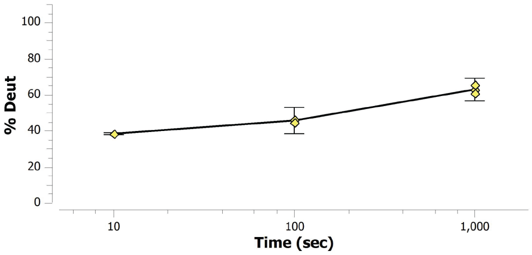

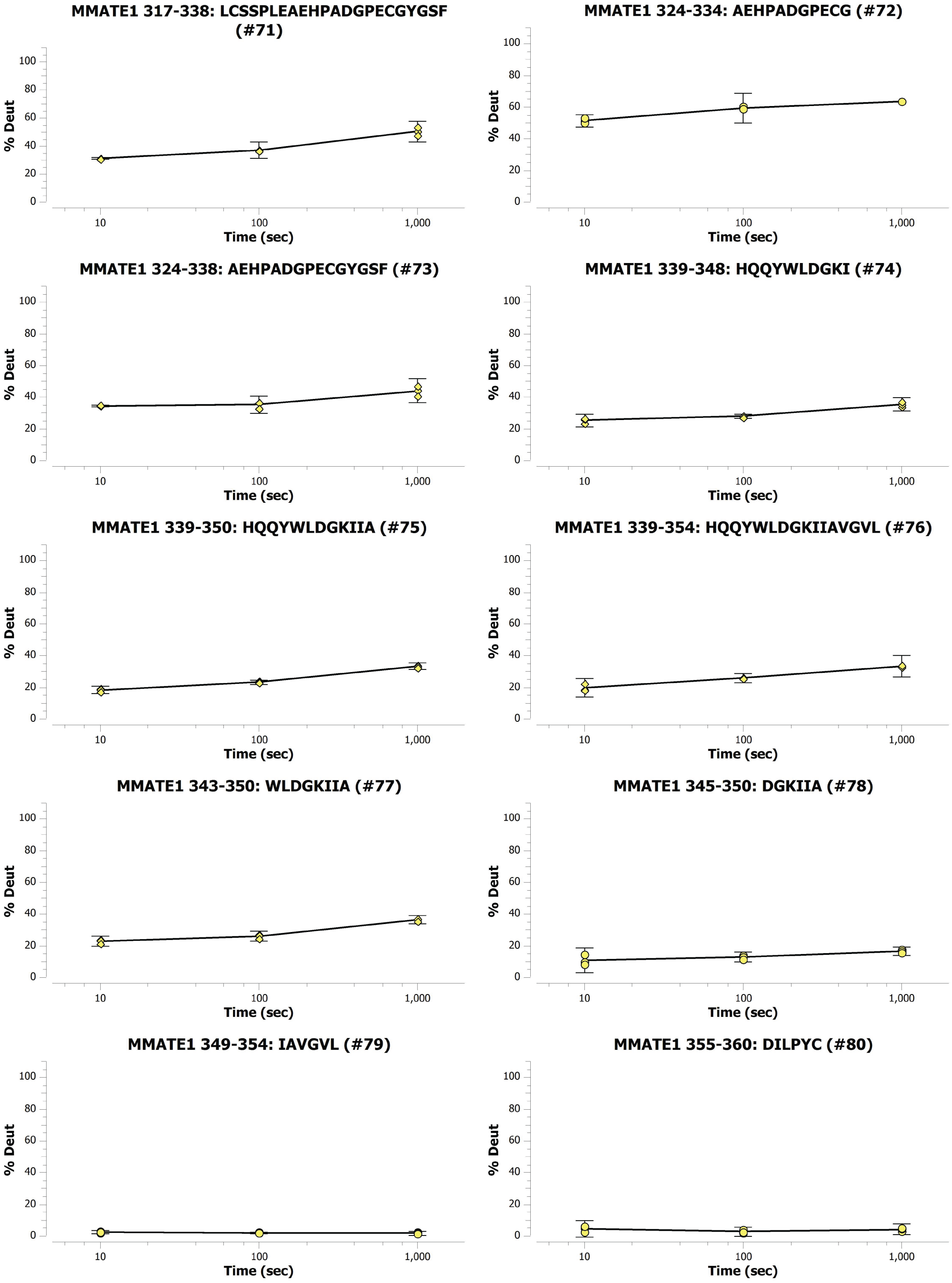
MMATE1 317-338: LCSSPLEAEHPADGPECGYGSF (#71)
MMATE1 324-334: AEHPADGPECG (#72)
MMATE1 324-338: AEHPADGPECGYGSF (#73)
MMATE1 339-348: HQQYWLDGKI (#74)
MMATE1 339-350: HQQYWLDGKIIA (#75)
MMATE1 339-354: HQQYWLDGKIIAVGVL (#76)
MMATE1 343-350: WLDGKIIA (#77)
MMATE1 345-350: DGKIIA (#78)
MMATE1 349-354: IAVGVL (#79)
MMATE1 355-360: DILPYC (#80)
% Deut
Time (sec)
0
20
40
60
80
100
10
100
1,000

**MMATE1 355-362: DILPYCVS (#81)**

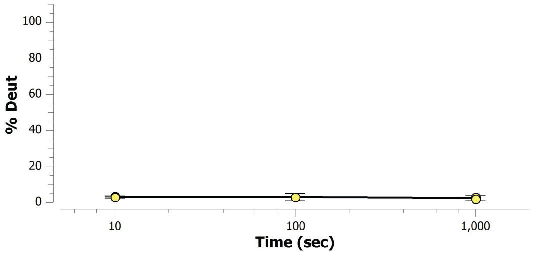


**MMATE1 355-363: DILPYCVSS (#82)**

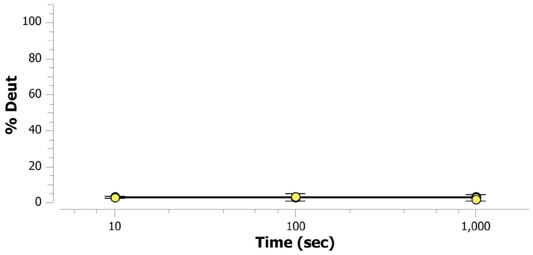


**MMATE1 355-364: DILPYCVSSV (#83)**

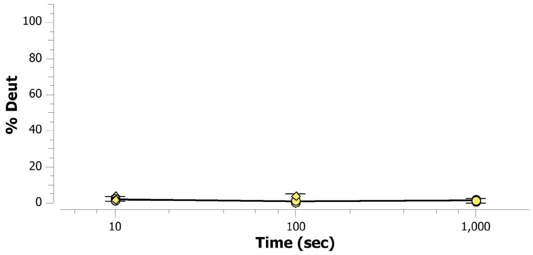


**MMATE1 357-369: LPYCVSSVYLYYD (#84)**

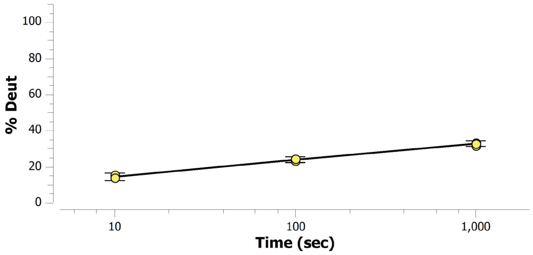


**MMATE1 365-374: YLYYDPDYSF (#85)**

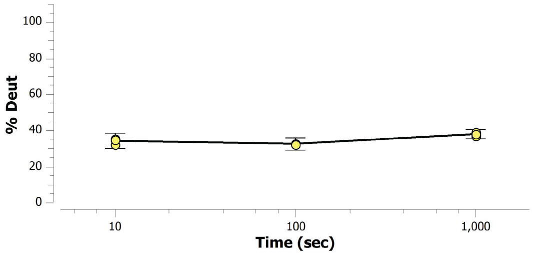


**MMATE1 366-374: LYYDPDYSF (#86)**

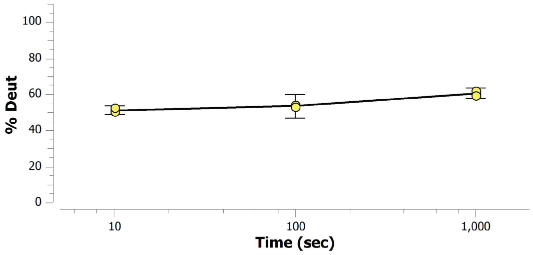


**MMATE1 367-374: YYDPDYSF (#87)**

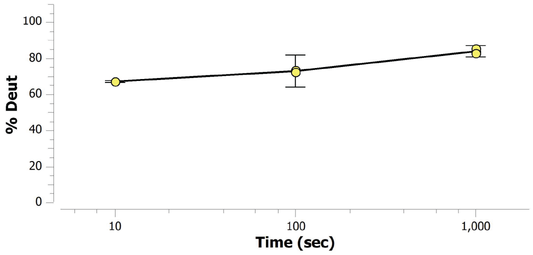


**MMATE1 367-375: YYDPDYSFL (#88)**

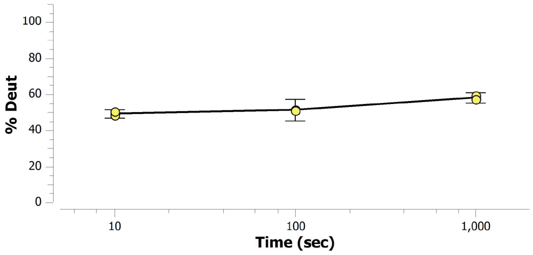


**MMATE1 375-380: LSLGVY (#89)**

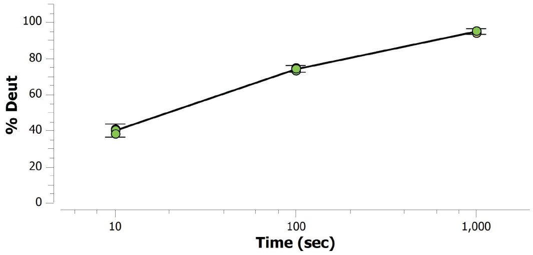


**MMATE1 375-382: LSLGVYSA (#90)**

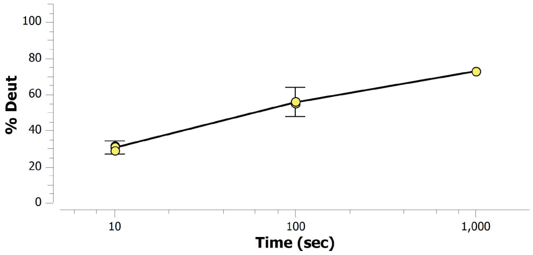

**MMATE1 386-401: IAFTRQLHEKTSQLSY (#91)**

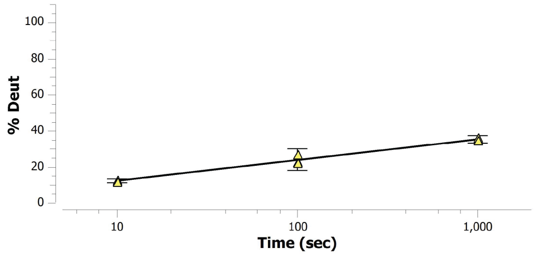


**MMATE1 389-398: TRQLHEKTSQ (#92)**

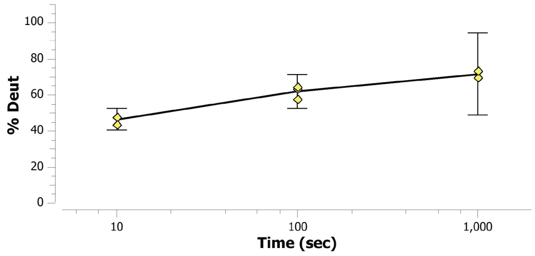


**MMATE1 389-399: TRQLHEKTSQL (#93)**

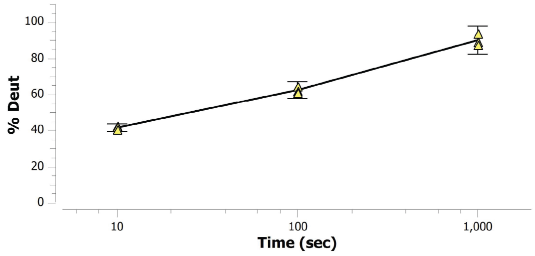


**MMATE1 389-401: TRQLHEKTSQLSY (#94)**

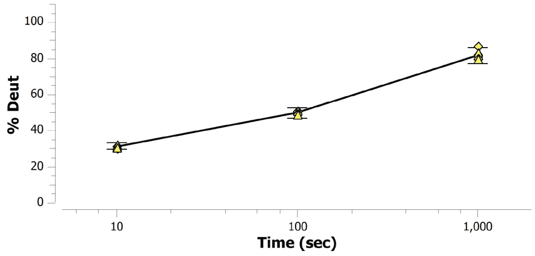


**MMATE1 400-406: SYYYMGF (#95)**

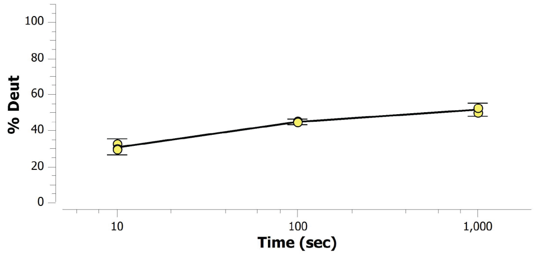


**MMATE1 407-425: YIHSCPKMRYKGQYIPSDL (#96)**

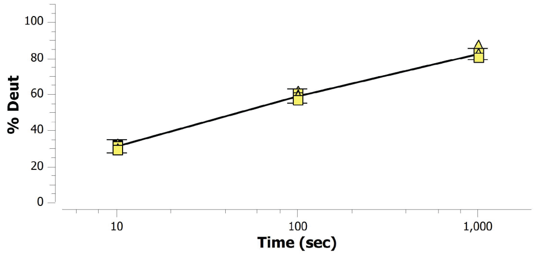


**MMATE1 407-426: YIHSCPKMRYKGQYIPSDLL (#97)**

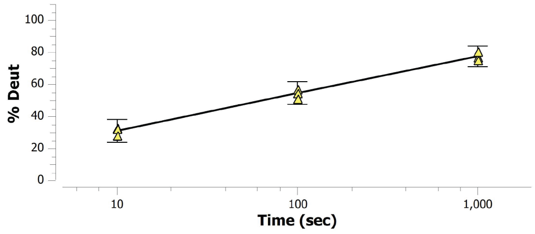


**MMATE1 407-427: YIHSCPKMRYKGQYIPSDLLC (#98)**

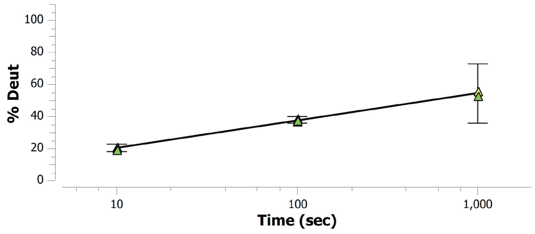


**MMATE1 415-425: RYKGQYIPSDL (#99)**

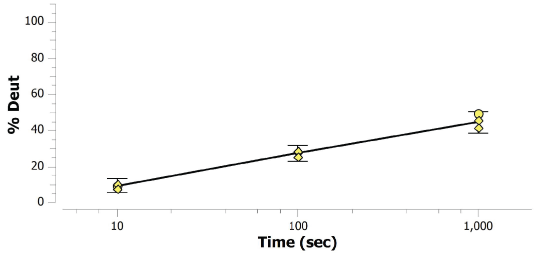


**MMATE1 423-431: SDLLCPETY (#100)**

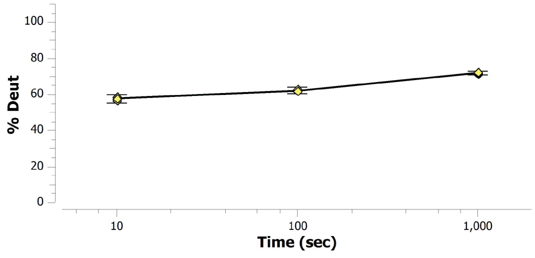

MMATE1 428-437: PETYVWVPIE (#101)

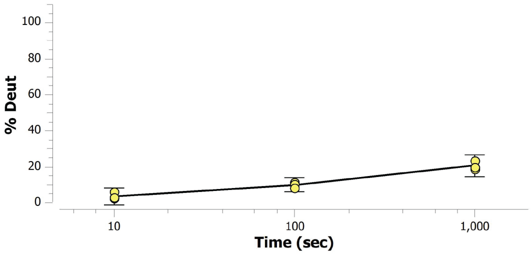


MMATE1 431-437: YVWVPIE (#102)

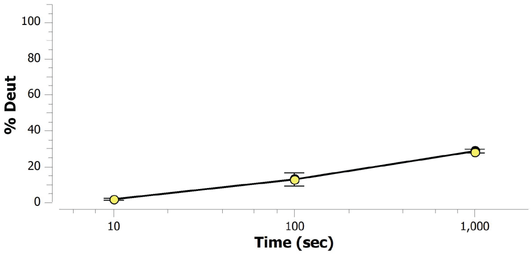


MMATE1 431-438: YVWVPIEQ (#103)

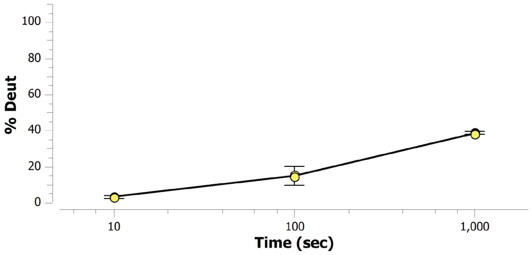


MMATE1 431-440: YVWVPIEQCL (#104)

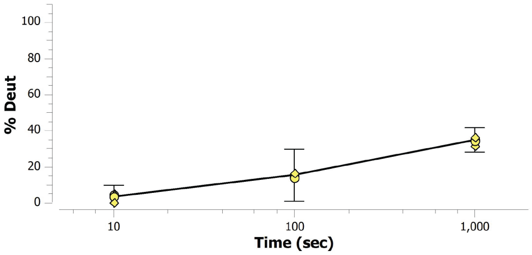


MMATE1 431-448: YVWVPIEQCLPSLDNSKY (#105)

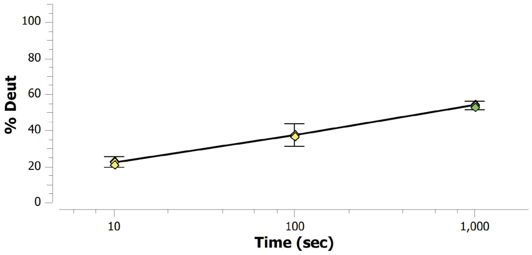


MMATE1 432-437: VWVPIE (#106)

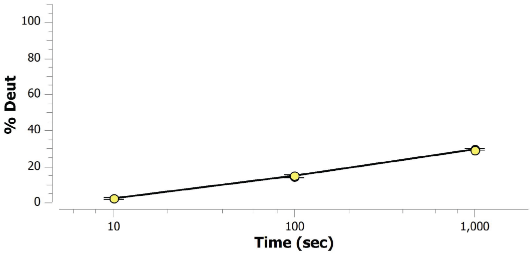


MMATE1 432-438: VWVPIEQ (#107)

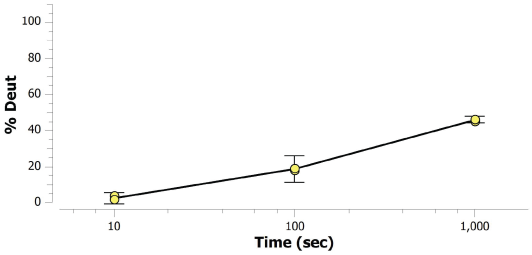


MMATE1 432-440: VWVPIEQCL (#108)

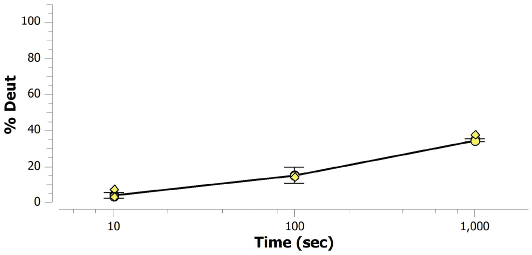


MMATE1 432-448: VWVPIEQCLPSLDNSKY (#109)

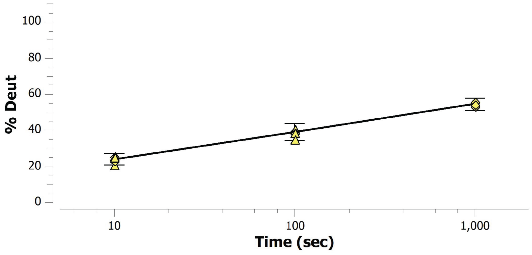


MMATE1 433-440: WVPIEQCL (#110)

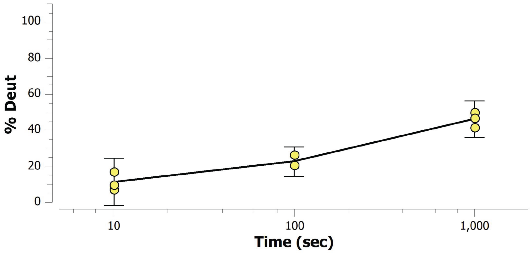

MMATE1 435-440: PIEQCL (#111)

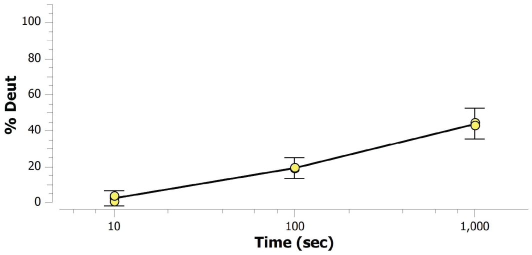


MMATE1 438-451: QCLPSLDNSKYCRF (#112)

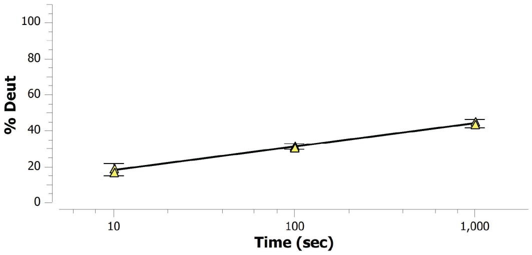


MMATE1 441-450: PSLDNSKYCR (#113)

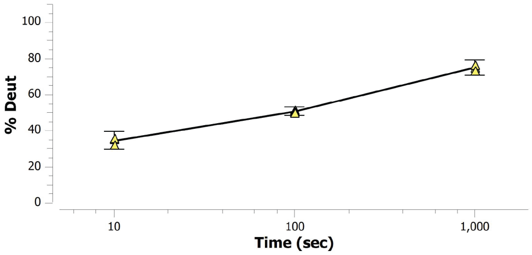


MMATE1 441-451: PSLDNSKYCRF (#114)

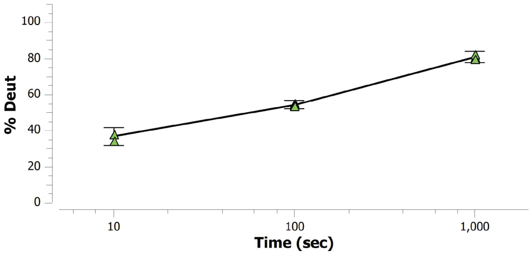


MMATE1 452-465: NQDPEAEDEGRSKE (#115)

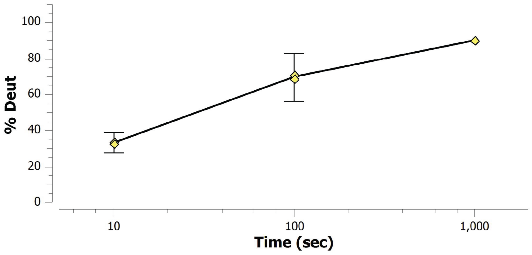


MMATE1 452-466: NQDPEAEDEGRSKEL (#116)

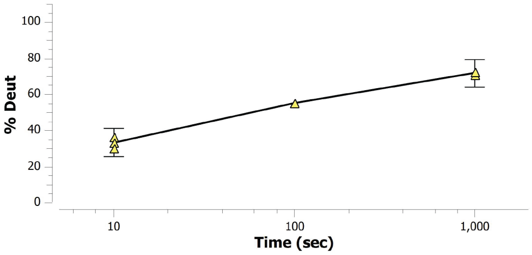


MMATE1 452-469: NQDPEAEDEGRSKELDRL (#117)

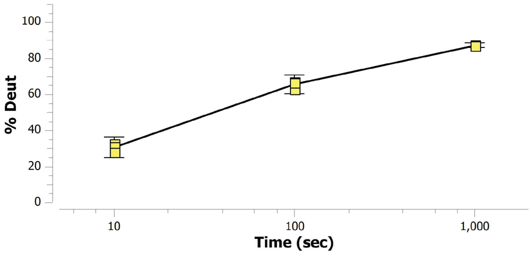


MMATE1 470-493:
RVFHRRSAMPYGVYKNHQEDPSEE (#118)

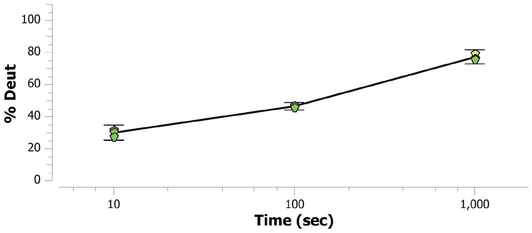


MMATE1 470-497:
RVFHRRSAMPYGVYKNHQEDPSEEAGVL (#119)

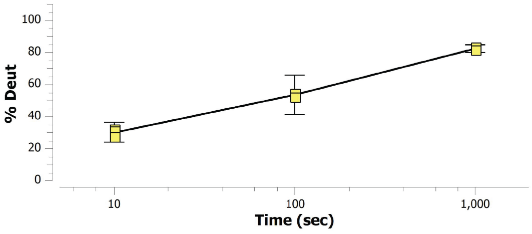


MMATE1 483-492: YKNHQEDPSE (#120)

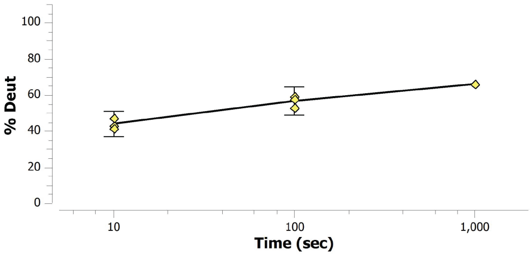

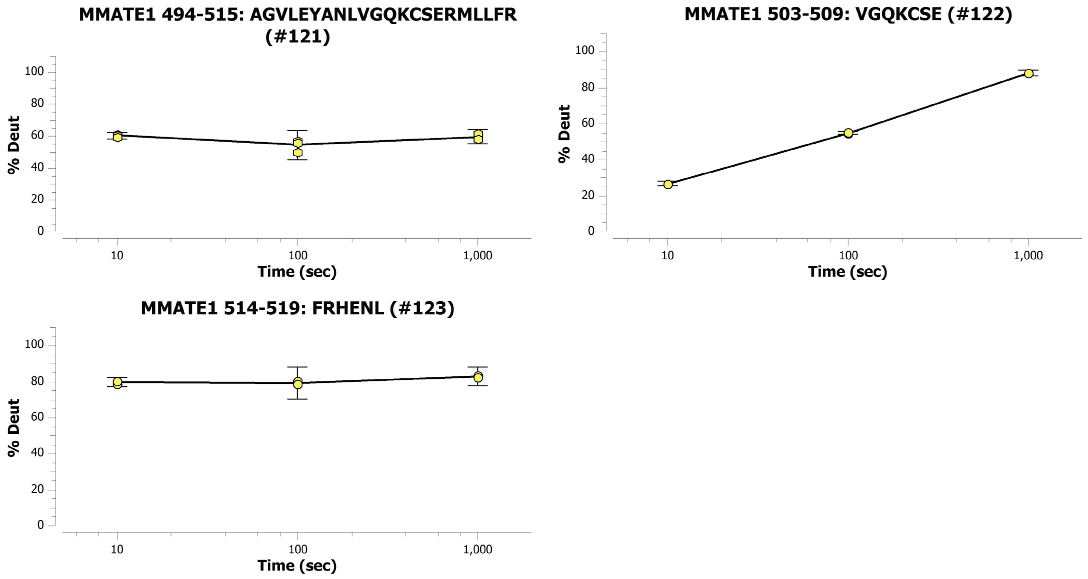


**Figure S9**. Kinetic percent deuteration plots of *Mm*ATE1-1 for all peptides at three time points.

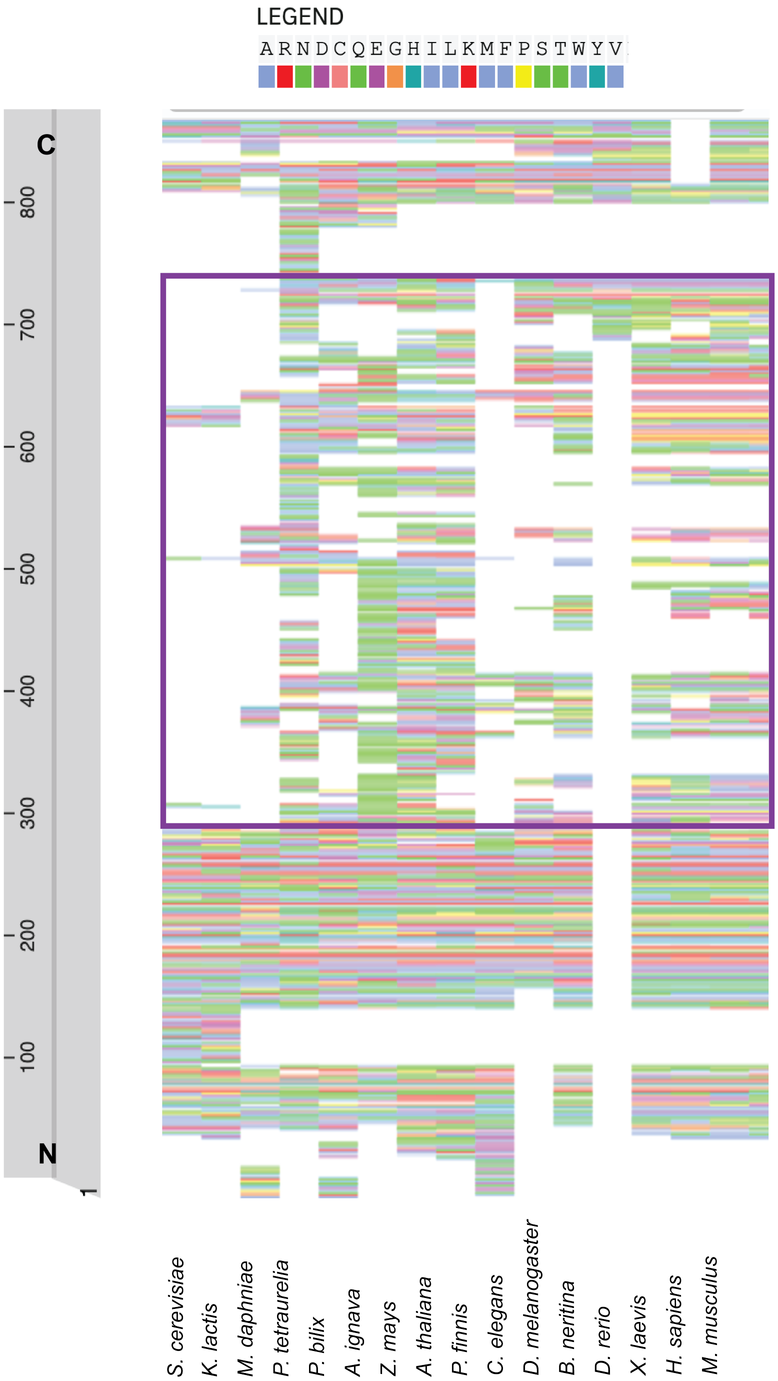


**Figure S10**. Large-scale MSA of ATE1 sequences illustrates how many, but not all, ATE1 sequences predict the presence of an IDR. The region that contains the IDR is highlighted with a purple box. The approximate locations of ATE1 sequences from key species are located to the left of the MSA. Notably missing the IDR region include yeast ATE1s (such as *S. cerevisiae* and *K. lactis*), while a shorter IDR is predicted in some common model organisms such as *C. elegans*.

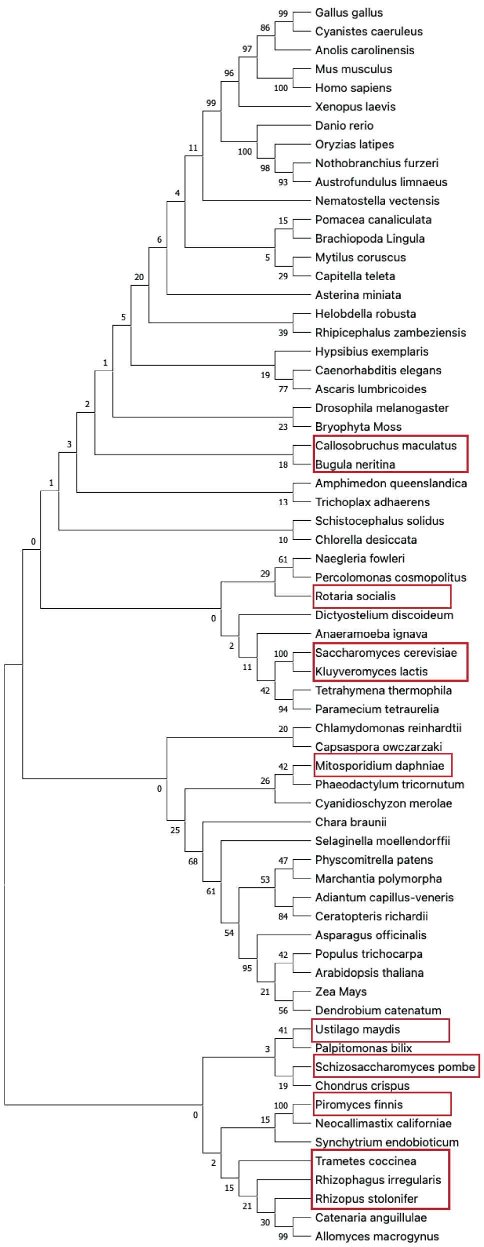


**Figure S11**. Phylogenic analysis of select ATE1 sequences. Organisms outlined with red boxes are predicted to be missing an IDR based on their lack of a low hydrophobicity but high overall net charge sequence within the middle of the predicted ATE1 protein. Notable is the lack of any clear divergence in the sequences with and without an IDR. In general, most (but not all) of the sequences predicted to lack an IDR are in the fungal kingdom, but several sequences are also predicted to have shorter IDRs than others (see Figure S13).

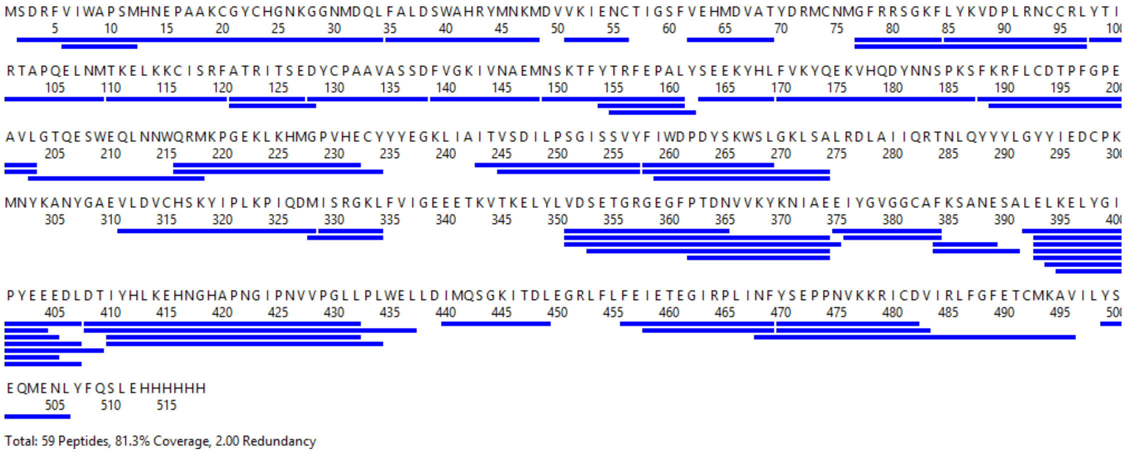


**Figure S12**. Sequence coverage map for *Sc*ATE1. Solid blue lines denote the peptic fragments analyzed in this study with total sequence coverage of 81.3%.

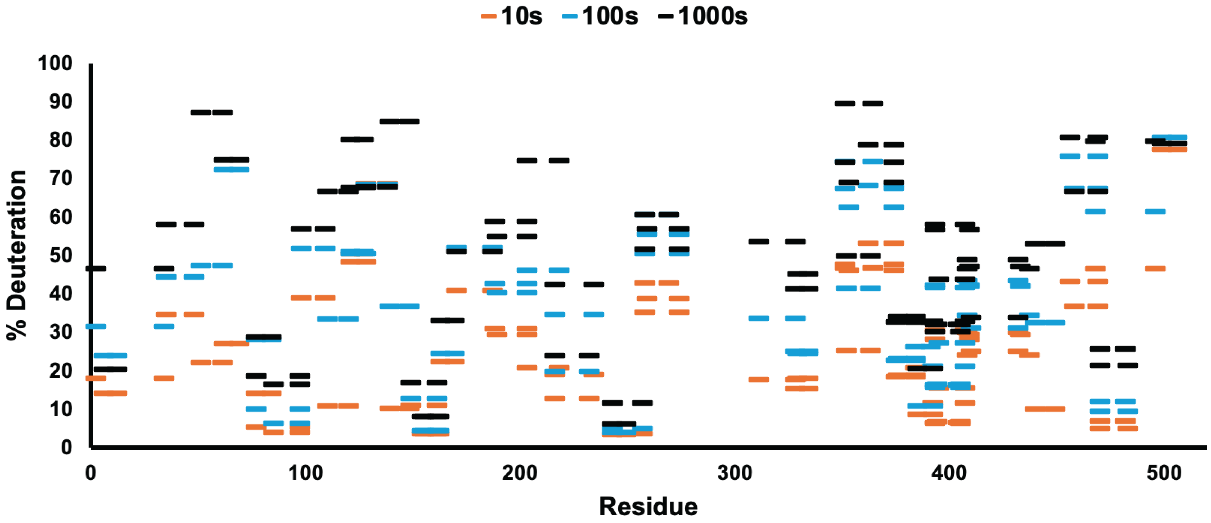


**Figure S13**. Woods plot of *Sc*ATE1-1 representing percent deuteration values in the apo state for time points 10s (yellow), 100s (blue), and 1000s (black), plotted against amino acid sequence.

**SCATE1 2-34: SDRFVIWAPSMHNEPAAKCGYCHGNKGGNMDQL (#1)**

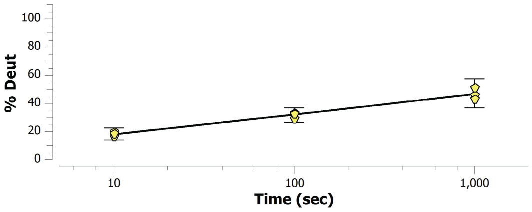


**SCATE1 6-12: VIWAPSM (#2)**

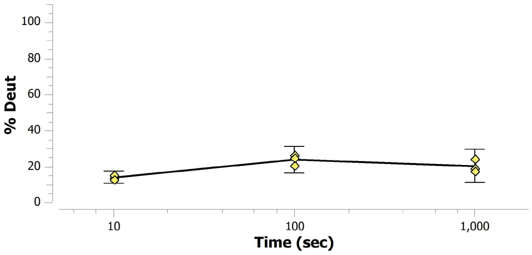


**SCATE1 35-48: FALDSWAHRYMNKM (#3)**

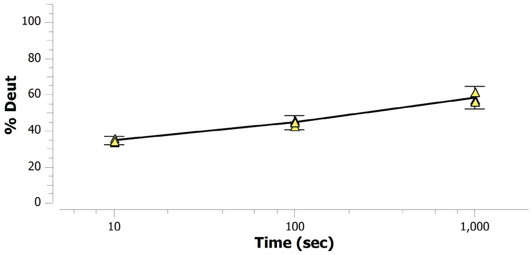


**SCATE1 51-61: VKIENCTIGSF (#4)**

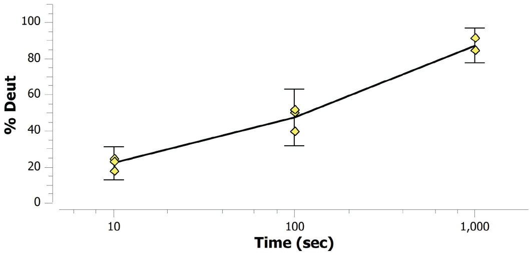


**SCATE1 62-69: VEHMDVAT (#5)**

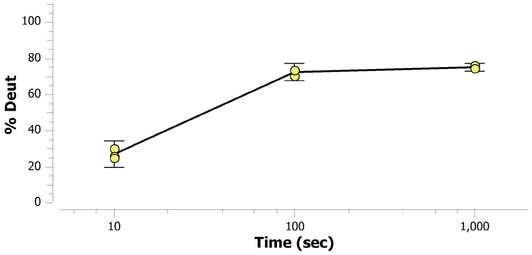


**SCATE1 77-84: GFRRSGKF (#6)**

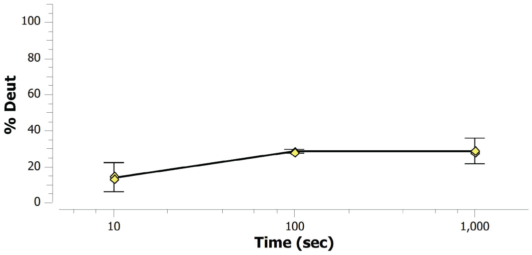


**SCATE1 77-97: GFRRSGKFLYKVDPLRNCCRL (#7)**

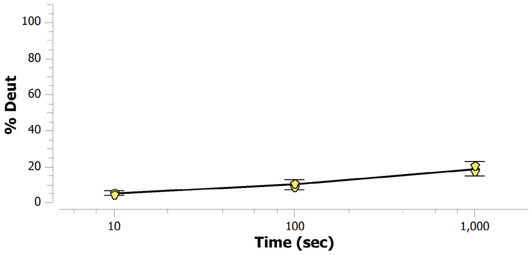


**SCATE1 85-97: LYKVDPLRNCCRL (#8)**

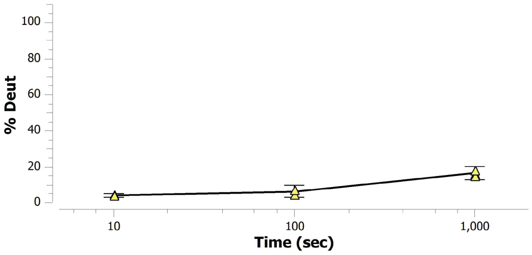


**SCATE1 98-109: YTIRTAPQELNM (#9)**

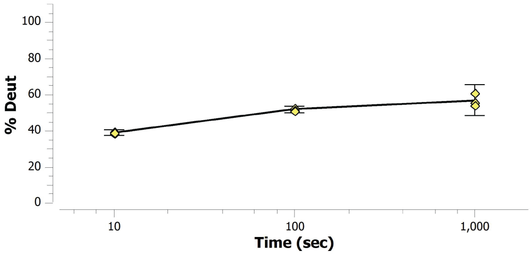


**SCATE1 110-120: TKELKKCISRF (#10)**

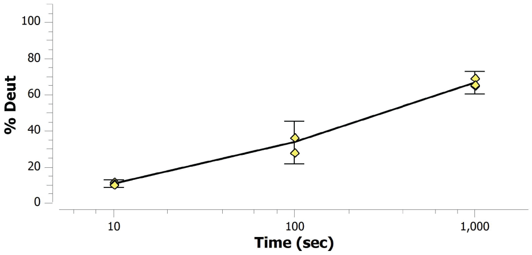

**SCATE1 121-127: ATRITSE (#11)**

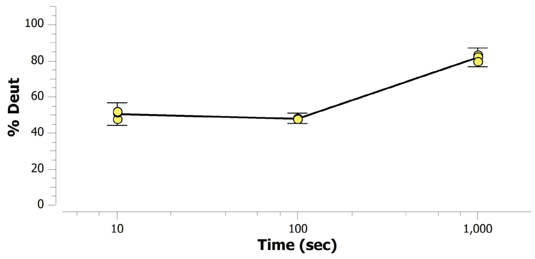


**SCATE1 121-128: ATRITSED (#12)**

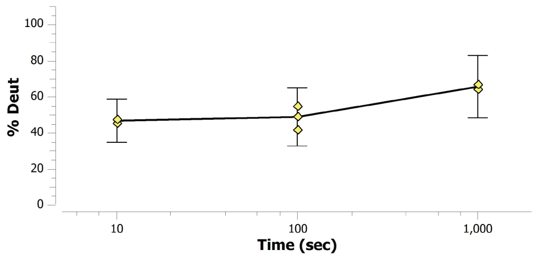


**SCATE1 128-138: DYCPAAVASSD (#13)**

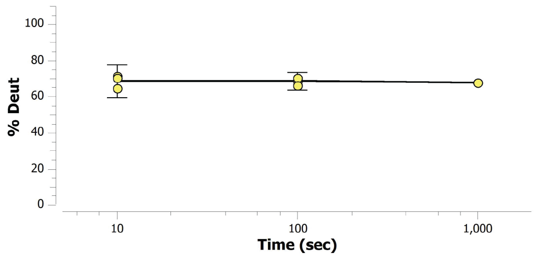


**SCATE1 139-148: FVGKIVNAEM (#14)**

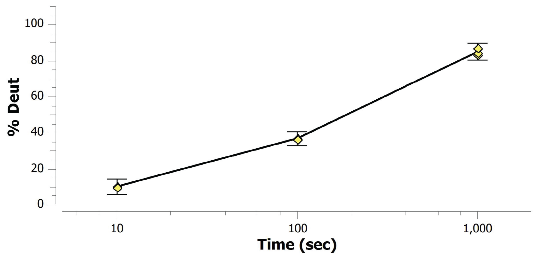


**SCATE1 149-161: NSKTFYTRFEPAL (#15)**

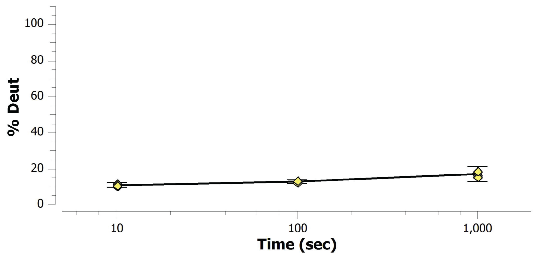


**SCATE1 154-161: YTRFEPAL (#16)**

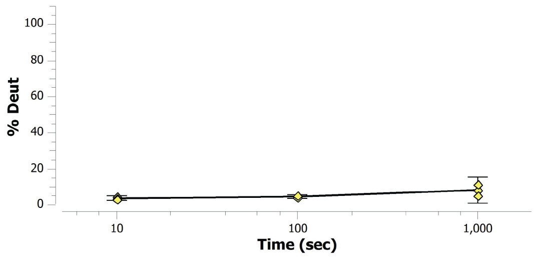


**SCATE1 155-162: TRFEPALY (#17)**

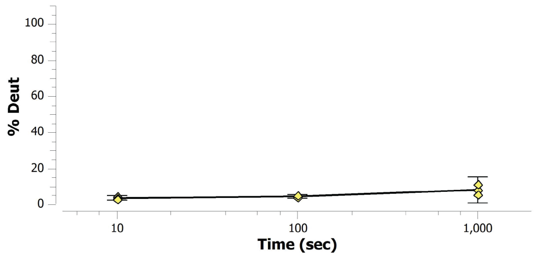


**SCATE1 163-169: SEEKYHL (#18)**

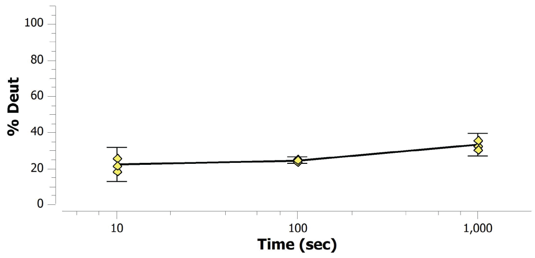


**SCATE1 170-187: FVKYQEKVHQDYNNSPKS (#19)**

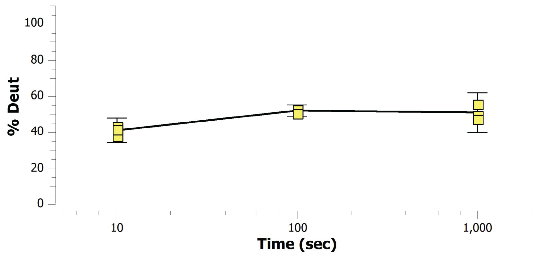


**SCATE1 188-203: FKRFLCDTPFGPEAVL (#20)**

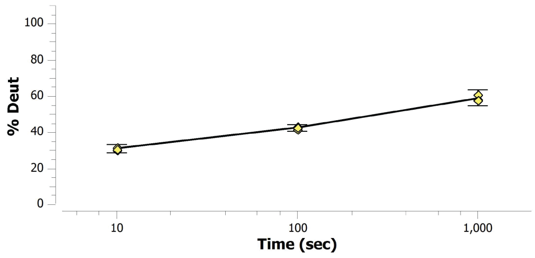

**SCATE1 189-203: KRFLCDTPFGPEAVL (#21)**

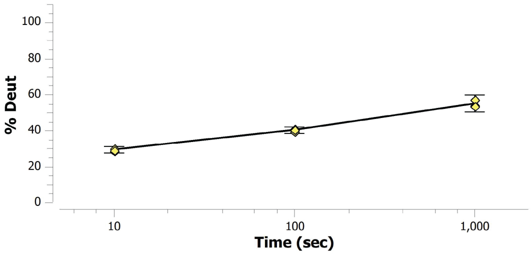


**SCATE1 203-218: LGTQESWEQLNNWQRM (#22)**

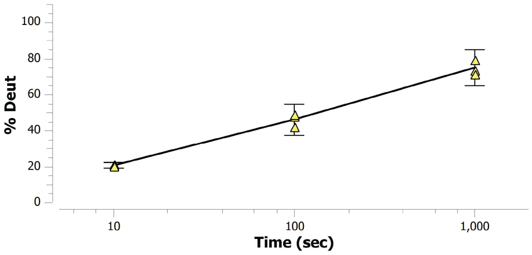


**SCATE1 216-232: QRMKPGEKLKHMGPVHE (#23)**

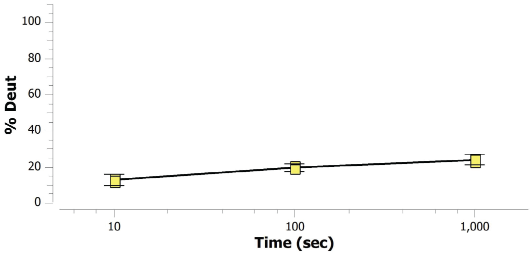


**SCATE1 216-234: QRMKPGEKLKHMGPVHECY (#24)**

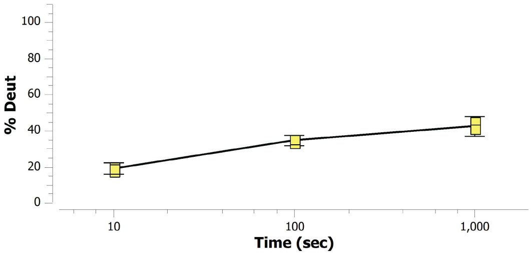


**SCATE1 243-249: ITVSDIL (#25)**

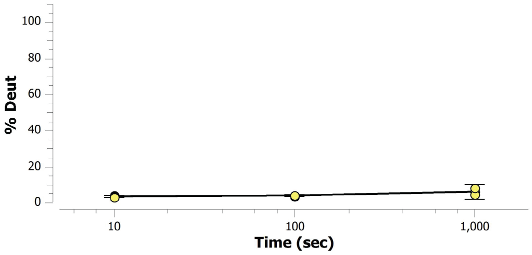


**SCATE1 243-257: ITVSDILPSGISSVY (#26)**

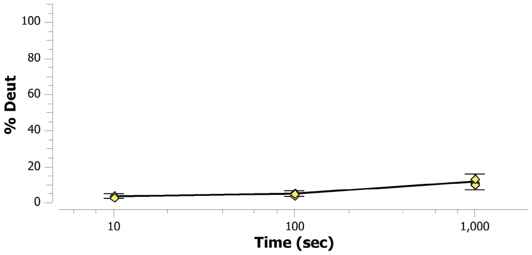


**SCATE1 258-269: FIWDPDYSKWSL (#27)**

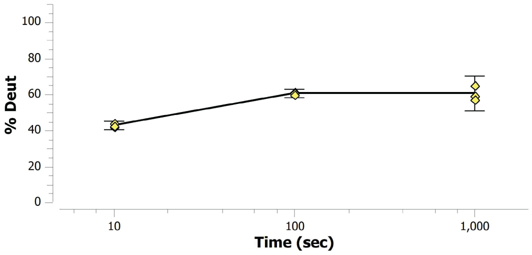


**SCATE1 258-274: FIWDPDYSKWSLGKLSA (#28)**

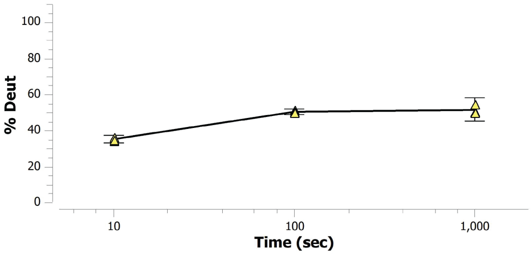


**SCATE1 259-274: IWDPDYSKWSLGKLSA (#29)**

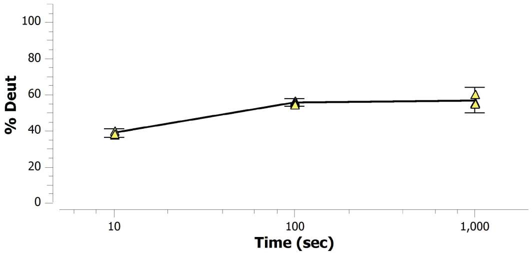


**SCATE1 311-328: VLDVCHSKYIPLKPIQDM (#30)**

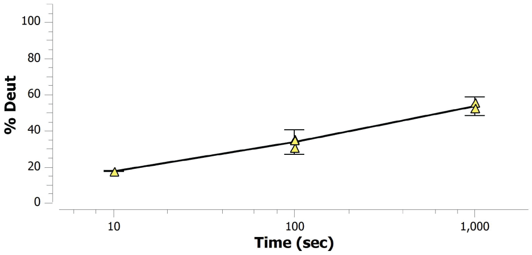

**SCATE1 328-334: MISRGKL (#31)**

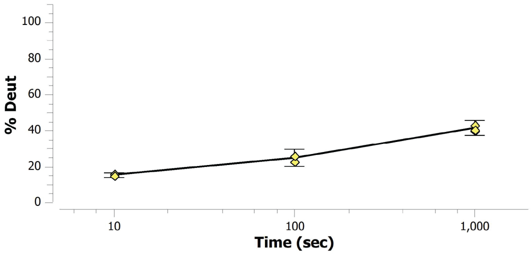


**SCATE1 329-334: ISRGKL (#32)**

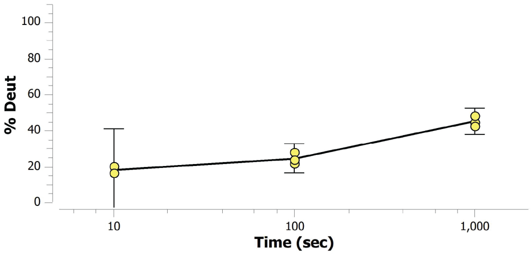


**SCATE1 351-364: VDSETGRGEGFPTD (#33)**

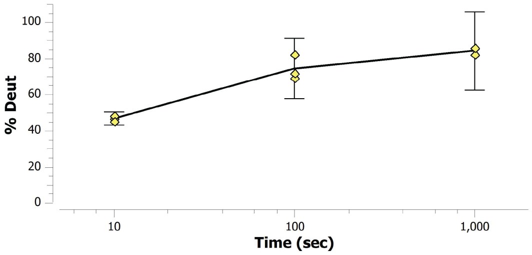


**SCATE1 351-374: VDSETGRGEGFPTDNVVKYKNIAE (#34)**

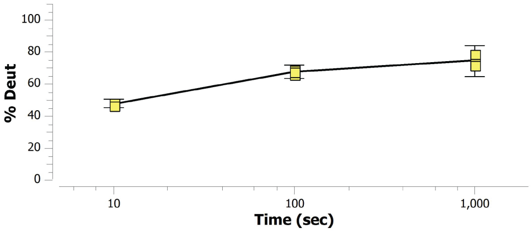


**SCATE1 352-363: DSETGRGEGFPT (#35)**

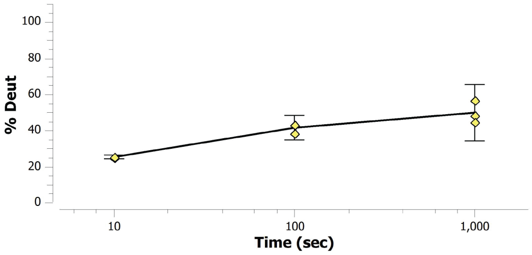


**SCATE1 353-374: SETGRGEGFPTDNVVKYKNIAE (#36)**

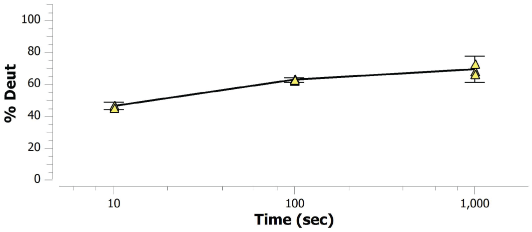


**SCATE1 362-374: PTDNVVKYKNIAE (#37)**

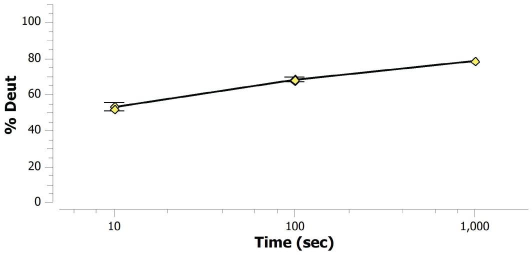


**SCATE1 375-384: EIYGVGGCAF (#38)**

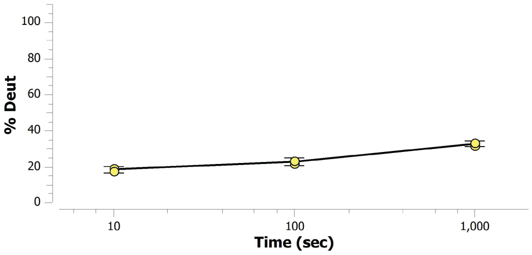


**SCATE1 376-384: IYGVGGCAF (#39)**

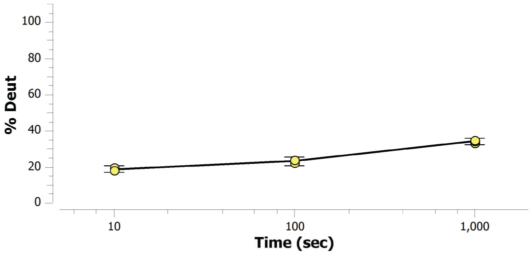


**SCATE1 384-391: FKSANESA (#40)**

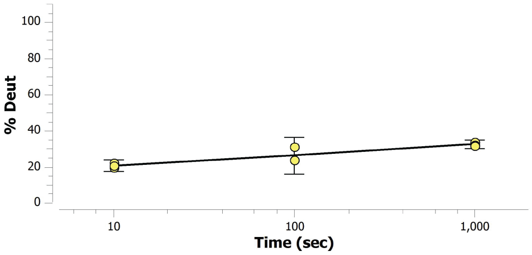

**SCATE1 385-392: KSANESAL (#41)**

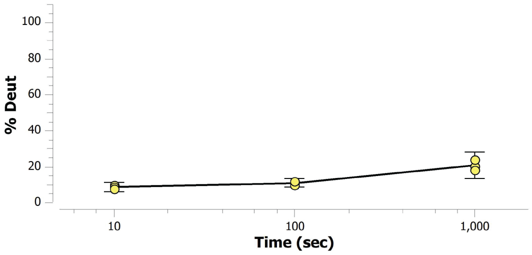


**SCATE1 392-407: LELKELYGIPYEEEDL (#42)**

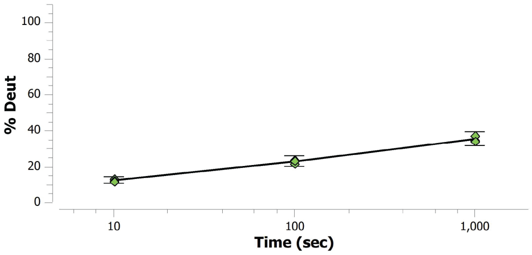


**SCATE1 393-404: ELKELYGIPYEE (#43)**

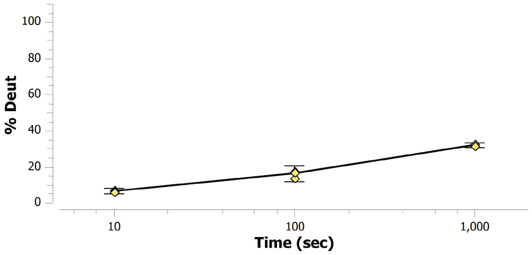


**SCATE1 393-405: ELKELYGIPYEEE (#44)**

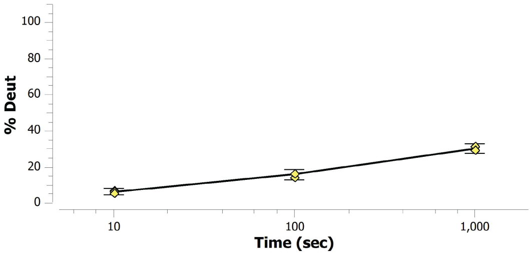


**SCATE1 393-407: ELKELYGIPYEEEDL (#45)**

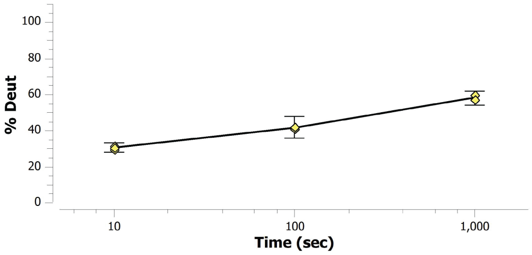


**SCATE1 393-409: ELKELYGIPYEEEDLDT (#46)**

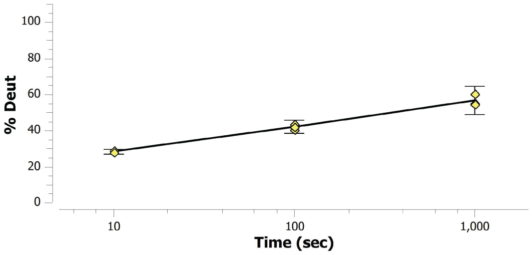


**SCATE1 394-405: LKELYGIPYEEE (#47)**

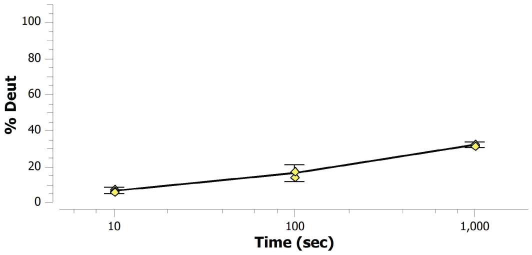


**SCATE1 395-407: KELYGIPYEEEDL (#48)**

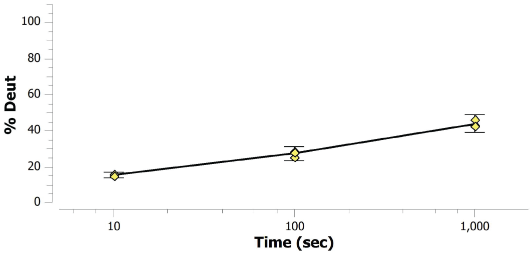


**SCATE1 408-432: DTIYHLKEHNGHAPNGIPNVVPGLL (#49)**

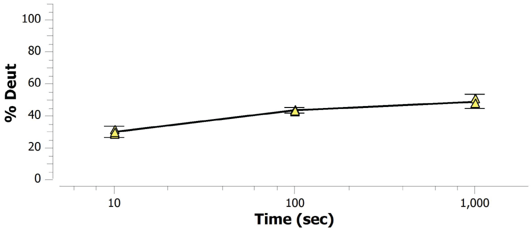


**SCATE1 408-437: DTIYHLKEHNGHAPNGIPNVVPGLLPLWEL (#50)**

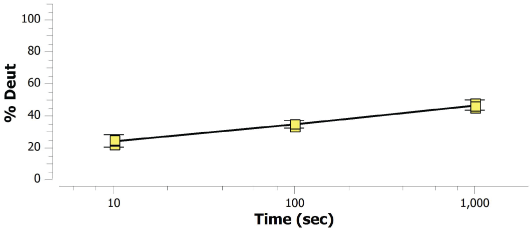

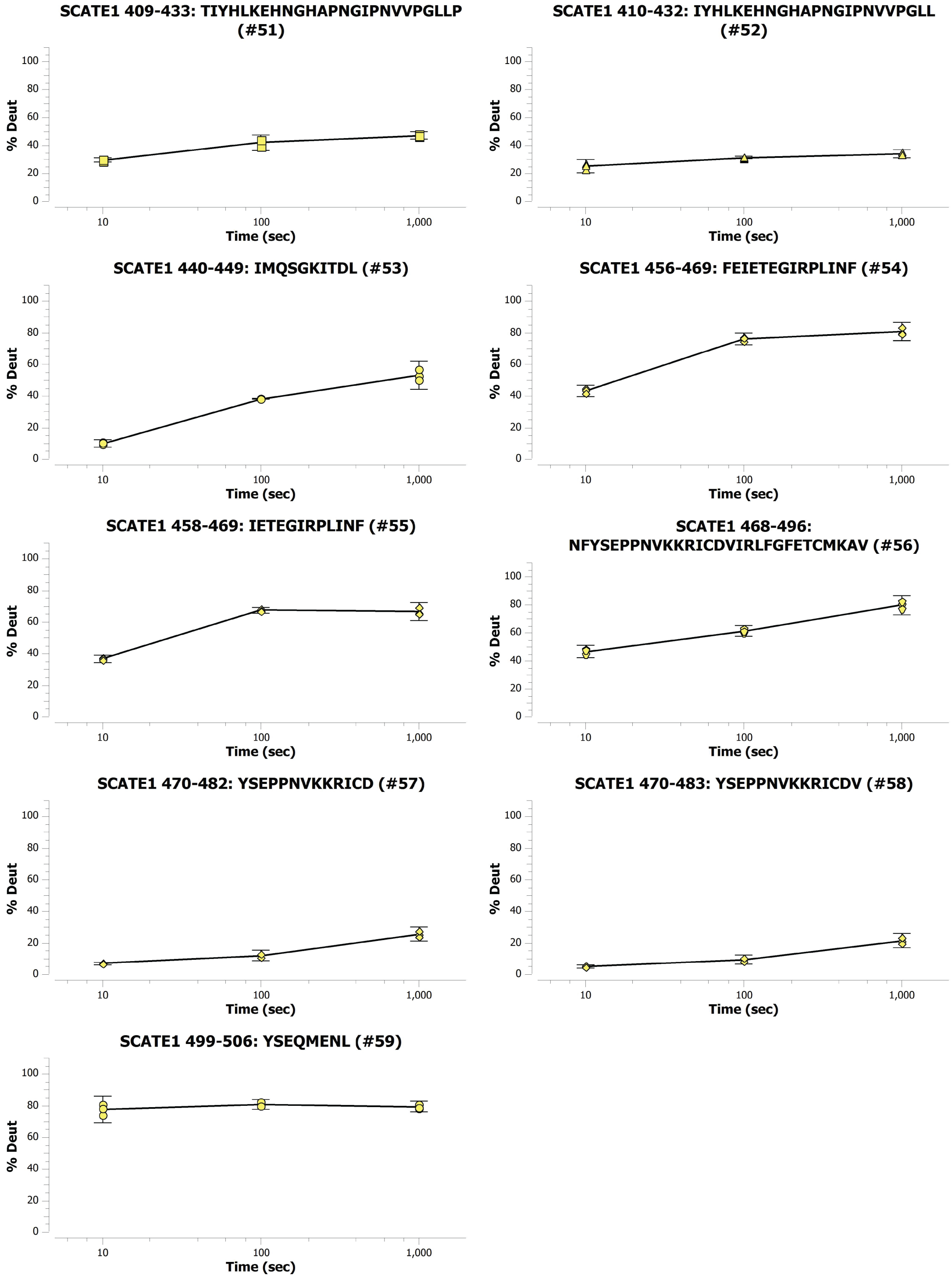


**Figure S14**. Kinetic percent deuteration plots of *Sc*ATE1 for all peptides at three time points.
